# Organometallic Pillarplexes that bind DNA 4-way Holliday Junctions and Forks

**DOI:** 10.1101/2023.01.04.522759

**Authors:** James S. Craig, Larry Melidis, Hugo D. Williams, Samuel J. Dettmer, Alexandra A. Heidecker, Philipp J. Altmann, Shengyang Guan, Callum Campbell, Douglas F. Browning, Roland K.O. Sigel, Silke Johannsen, Ross T. Egan, Brech Aikman, Angela Casini, Alexander Pöthig, Michael J. Hannon

**Affiliations:** Physical Sciences for Health Centre, University of Birmingham, Edgbaston, Birmingham B15 2TT, UK; School of Chemistry, University of Birmingham, Edgbaston, Birmingham B15 2TT, UK; Department of Chemistry & Catalysis Research Center, Technical University of Munich, Lichtenbergstr. 4, 85748 Garching, Germany; School of Biosciences, University of Birmingham, Edgbaston, Birmingham B15 2TT, UK; Department of Chemistry, University of Zürich, Winterthurerstr. 190, 8057 Zürich, Switzerland; Chair of Medicinal and Bioinorganic Chemistry, Department of Chemistry, Technical University of Munich (TUM), Lichtenbergstr. 4, 85748 Garching b. München, Germany

**Keywords:** Supramolecular, DNA, Four-way junctions, Holliday junction, Bioinorganic, Pillarplex, Y-shaped forks

## Abstract

Holliday 4-way junctions are key to important biological DNA processes (insertion, recombination and repair) and are dynamic structures which adopt either open or closed conformations, with the open conformation being the biologically active form. Tetracationic metallo-supramolecular pillarplexes display aryl faces about a cylindrical core giving them an ideal structure to interact with the central cavities of open DNA junctions. Combining experimental studies and MD simulations we show that an Au pillarplex can bind DNA 4-way junctions (Holliday junctions) in their open form, a binding mode not accessed by synthetic agents before. The Au pillarplexes can bind designed 3-way junctions too but their large size leads them to open up and expand that junction, disrupting the base pairing which manifests in an increase in hydrodynamic size and a lower junction thermal stability. At high loading they re-arrange both 4-way and 3-way junctions into Y-shaped DNA forks to increase the available junction-like binding sites. The structurally related Ag pillarplexes show similar DNA junction binding behaviour, but a lower solution stability. This pillarplex binding contrasts with (but complements) that of the metallo-supramolecular cylinders, which prefer 3-way junctions and we show can rearrange 4-way junctions into 3-way junction structures. The pillarplexes’ ability to bind open 4-way junctions creates exciting possibilities to modulate and switch such structures in biology, as well as in synthetic nucleic acid nanostructures where they are key interconnecting components. Studies in human cells, confirm that the pillarplexes do reach the nucleus, with antiproliferative activity at levels similar to those of cisplatin. The findings provide a new roadmap for targeting higher order junction structures using a metallo-supramolecular approach, as well as expanding the toolbox available to design bioactive junction-binders into organometallic chemistry.

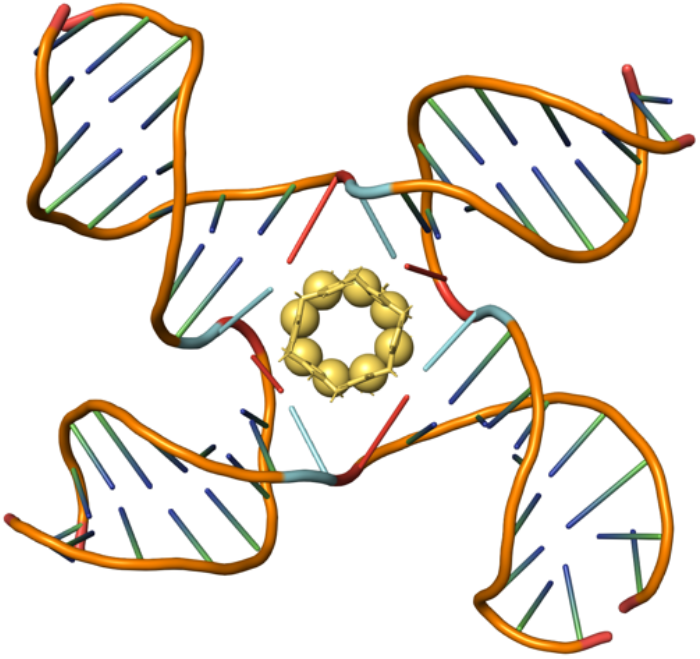

## INTRODUCTION

As the repository of our genetic code, DNA is an important and fascinating target. Agents that can bind to it and potentially regulate its processing have enormous potential. Indeed, synthetic agents that target the DNA duplex such as cisplatin and doxorubicin, are key agents in the clinic in the fight against many cancers.^1–4^ The duplex is the most common form of DNA in the body, but when the information is being accessed and processed a range of other structures are formed, notably the fork structures associated with DNA transactions.^5,6^ The importance of non-duplex structures in genomic DNA is increasingly recognised: Junctions such as the Holliday junction are involved in DNA repair and viral insertion,^7^ and G4-quadruplexes are implicated in regulation of some gene promoters as well as in telomere stability,^8– 12^ while the roles of other known cellular DNA structures such as i-motifs,^13–15^ B-Z junctions^16^ and threeway junctions as found at triplet repeat expansions,^17,18^ are still being elucidated. These are all attractive as targets that afford a level of selectivity in DNA binding – a structural selectivity that complements and expands traditional attempts at sequence selective recognition.

Among these non-canonical structures, the most explored has been the G4-quadruplex, with many elegant agents reported that offer large flat planar surfaces to interact with the ends of the quadruplex stack.^8,10–12,19–22^ Junctions are less well-studied,^23,24^ though we have shown that dinuclear metallo-supramolecular cylinders bind inside the heart of three-way junctions (3WJ) and characterised this binding by X-ray crystallography and NMR.^25–31^ The striking features of this binding are the cavity fit and the face-face pi-stacking the surface of the cylinder makes with the DNA-bases at the junction point. The key to this is that the aryl rings in the centre of the cylinder present their aryl faces to the outside of the structure to create pi-surfaces^25–28^ and contrasts with typical polypyridyl metal complexes (such as the archetypal ruthenium tris bipyridine or tris phenanthroline complexes) which present their CH lined aryl edges instead. Monchaud has screened libraries of agents for 3WJ binding and identified as his lead agent an organic 3-arm cryptand-type system with aryl rings that could potentially rotate to present a very similar aryl surface conformation to the cylinders.^32–35^ Although the binding is not yet structurally characterised, very recent MD simulations suggest this cryptand may also be able to thread into a 3WJ and bind as the cylinder does.^36^ Other 3WJ binders^32,34,35,37–46^ that lack these outward-facing pi-surfaces may bind outside the junction cavity.

Four-way or Holliday junctions (4WJ) are different in nature to 3WJ. While an open cruciform-style structure is possible and is seen in complex with proteins, in their absence the junction cavity closes up allowing the arms to come together in coaxial stacks forming a stacked X configuration. Searcey has taken advantage of this and designed binders that contain two linked intercalators to bind two adjacent arms and with Cardin has structurally characterised this binding mode.^47,48^ Bonnet^49^ has characterised a very interesting 4WJ-like discontinuous structure assembled from a hexanucleotide and containing a small metallo-intercalator inserting between inter-duplex pairs. Binding of organic intercalators in duplex arms has also been shown adjacent to the junction point in 4WJs.^50–52^ The open cruciform-style 4WJ junction cavity has not been addressed with synthetic binders.

Based on our analysis of key binding features, we sought other designs that also offer external pi-face surfaces and thus might be suited to junction binding, and identified another new class of cationic supramolecular organometallic complexes,^53^ called pillarplexes.^53,54^ These structures are composed of two organic macrocyclic ligands, each with 6 aryl (imidazole/pyrazole or triazole) rings, that linearly coordinate 8 gold or silver centres (Figure 1). The pillarplexes are a similar width to the cylinder but of a more circular girth (the junction binding unit of the cylinder is in fact a twisted triangular prism that could be circumscribed by the pillarplex - see SI Figure S1). The pillarplexes are significantly shorter than the cylinders (1.2 nm *c*.*f*. 1.9 nm), but importantly they have the same tetra-cationic charge that will contribute to the binding to anionic DNA. We show herein that these pillarplex structures can bind to open DNA 4-way junctions as well as 3-way junctions and Y-shaped forks. Since they incorporate N-heterocyclic carbene ligands, they also now extend design of nucleic acid junction binders into organometallic chemistry with the different design opportunities this offers for the future.

**Figure 1:**
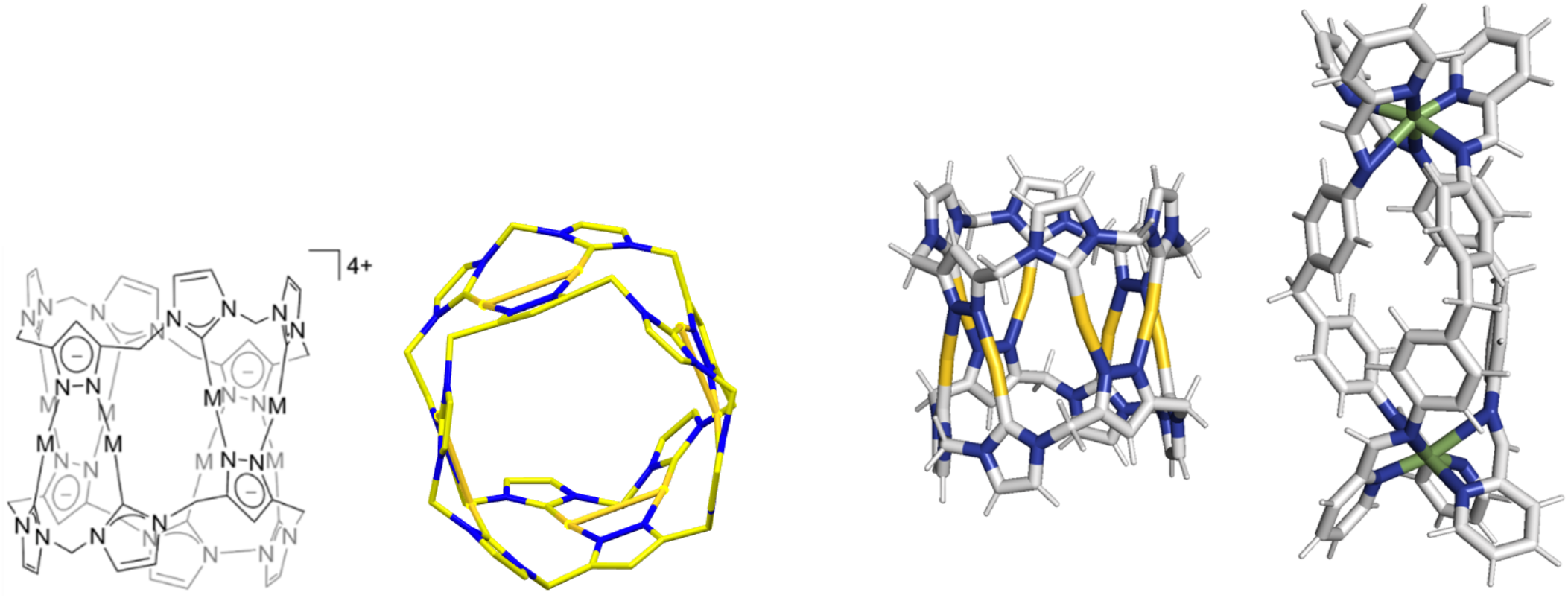
Left: structure of the pillarplex (M=Ag(I)/Au(I)) showing its chemical composition, and a view through its cylindrical axis emphasising its 4 fold symmetry. Right: comparison of the dimensions of the Au pillarplex and the Ni(II)/Fe(II) cylinders, that are compared in this work. See also Figure S1 for further comparative images.

## RESULTS AND DISCUSSION

Our initial experiment was to explore whether this pillarplex structure would bind with DNA three-way junctions (3WJ). We carried out a PAGE gel with three 14-mer strands whose sequences (Figure 2) are designed to assemble a perfect 3WJ.^29^ In this assay, the 3WJ is not formed at room temperature in absence of drugs and only single stranded DNA is observed (Figure 3 lane 1). If a drug binds in and stabilises the 3WJ then a new 3WJ band will be observed in the gel (as seen for cylinder used as positive control^25–28^ in lanes 11-14).

**Figure 2:**
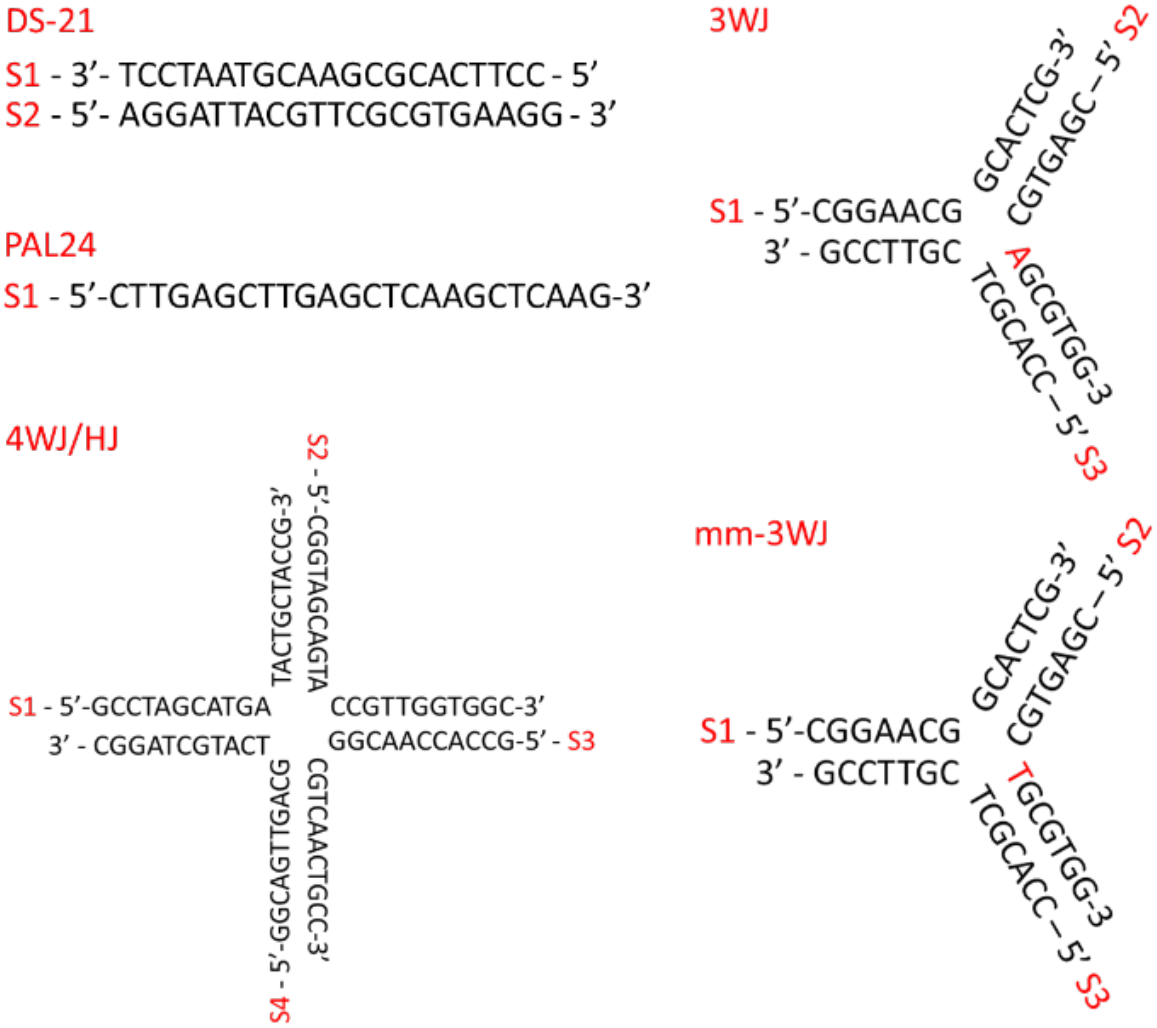
DNA sequences used in these studies.

**Figure 3.**
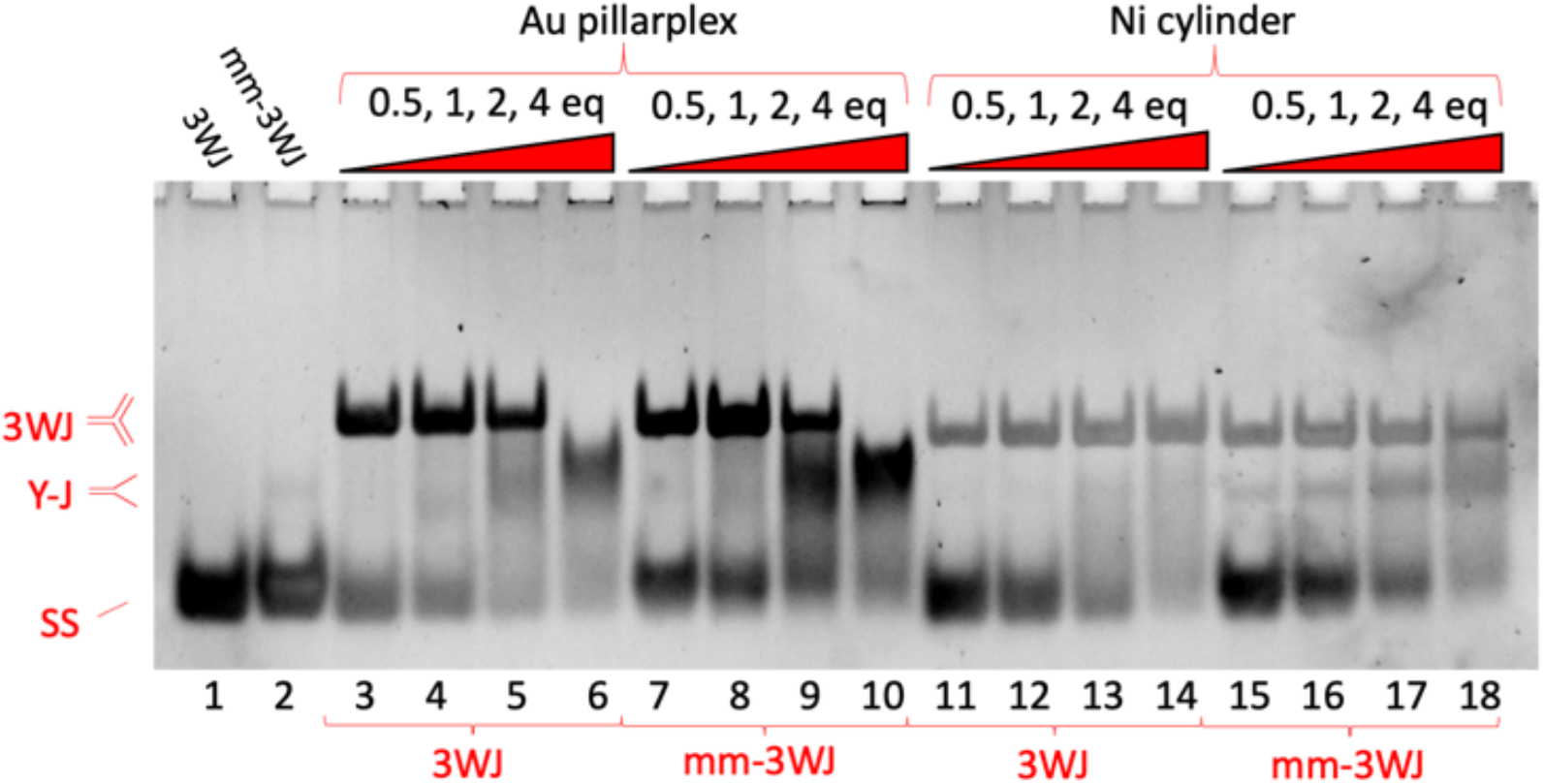
PAGE gel of two different DNA structures (3WJ and mm-3WJ) incubated with different complexes (Au pillarplex and Ni cylinder as positive control) at varying ratios (all gel lanes read from left to right). The 3WJ and mm-3WJ strand sequences used are shown in Figure 2. Controls (3 strands, no complexes) in lanes 1-2, then 3WJ is mixed with a complex at 0.5, 1, 2 and 4 eq (lanes 3-6 and 11-14), and mm-3WJ mixed with a complex at 0.5, 1, 2, 4 eq (lanes 7-10 and 15-18).

On addition of 0.5, 1 and 2 equivalents of the Au pillarplex (Figure 3 lanes 3-5), a 3WJ is formed, confirming that the Au pillarplex can bind the 3WJ. However, at four equivalents of pillarplex (lane 6) we see the 3WJ structure is replaced by a large, smeared band corresponding to 2-stranded Y-shaped fork junctions. This suggests that while the pillarplex prefers 3WJ over Y-shaped (or partially double stranded) forks, this preference is not strong and the pillarplex would rather bind forks than be unbound (the strands can form 50% more forks (2 strands) than perfect 3WJs (3 strands)). The broad nature of the Y-shaped band likely reflects the three different Y-structures that can be formed by the 3 strands and the flexibility of the single stranded branches. Controls with metal-free pillarplex ligand show no interactions with the DNA (Figure S2).

An interesting feature of the bound 3WJ band is that it has a slightly lower mobility in the gel compared to the 3WJ bound to the cylinder. This indicates that the 3WJ structure formed with the Au pillarplex has a larger hydrodynamic radius.

While the cylinder crystallises with a DNA hexamer,^26,27^ attempts to crystallise the pillarplex with the same DNA hexamer were unsuccessful, and the pillarplex precipitated the DNA at the concentrations needed for NMR experiments, preventing detailed structure characterisation. We, therefore, attempted to gain some atomistic insight using computational methods, using approaches previously used to explore cylinder and azacryptand 3WJ binding.^31,36,55,56^ As our starting point we used the crystal structure of the cylinder in complex with a 3WJ formed from a palindromic DNA hexamer^27^ (this structure is extremely similar to that of a 3WJ in complex with a protein^57^ and thus more broadly representative). On removing the bound cylinder, the unsupported 3WJ was not stable under molecular dynamics simulations, collapsing within 1 microsecond. For this reason we first used docking experiments to bring the pillarplex close to the 3WJ, where it sat outside the cavity, and then used this as the starting point for molecular dynamics simulations. The pillarplex reproducibly entered the cavity across multiple simulations, adopting a position with its pi surfaces facing the walls of the cavity (Figure 4) where it remained. However, in each case, this was associated with the breaking of a base pair at the junction point to enlarge the cavity and accommodate the pillarplex. The larger width dimensions/shape of the pillarplex (compared to the cylinders) is forcing the central cavity of the 3WJ to expand, which is consistent with our experimental observation of an increased hydrodynamic radius in the gel studies. By contrast in our simulations of the cylinder with the 3WJ DNA the cylinder resided in the cavity with no disruption to the base pairing even over long simulations (>10 ms).

**Figure 4.**
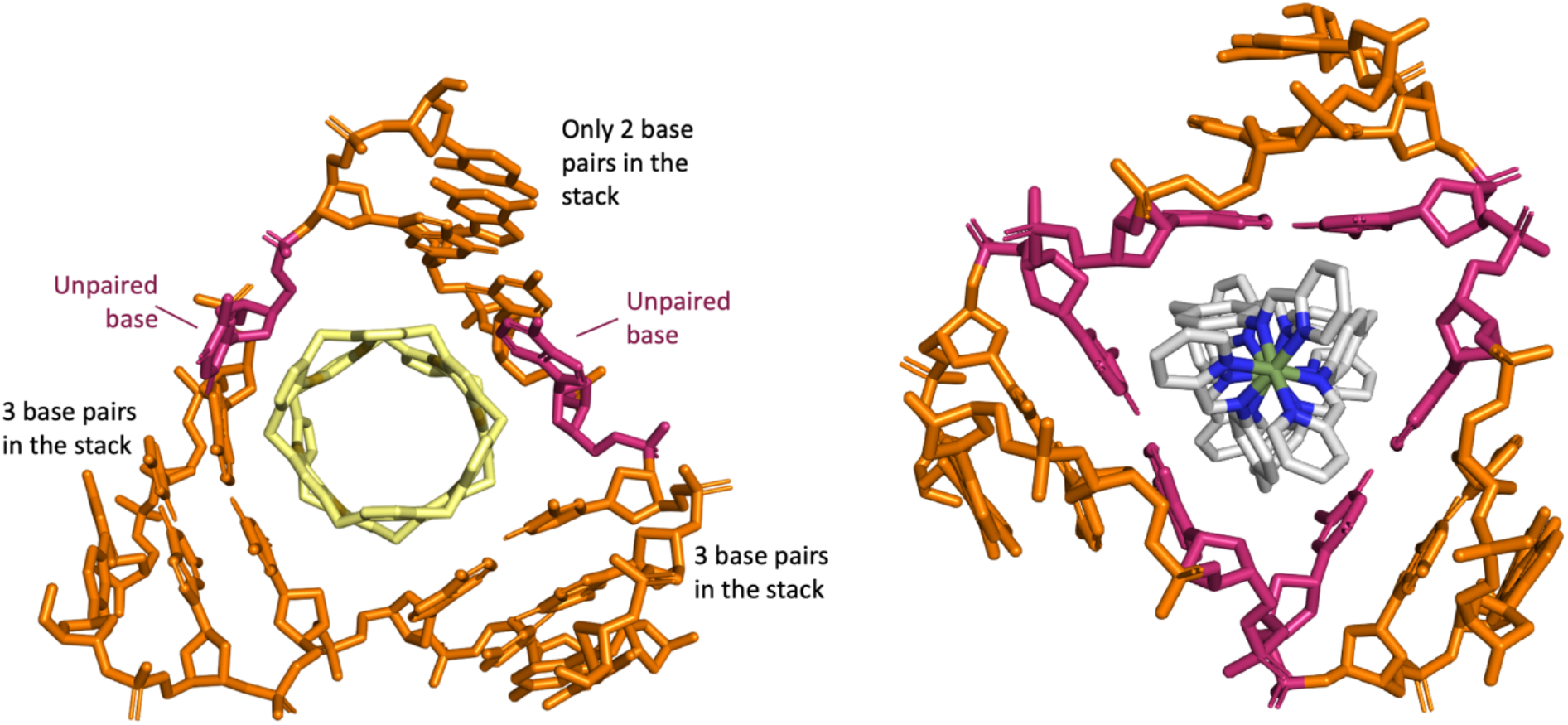
Left: MD simulations of Au pillarplex with palindromic DNA hexamer oligo (from pdb 3I1D). The MD shows Au Pillarplex inserting into the central cavity of the 3WJ where it opens up a base pair (highlighted in pink) down one arm thereby expanding the 3WJ cavity. Right: MD simulation of the Fe cylinder in the 3WJ where the central bases (pink) remain paired throughout the simulations, as observed experimentally in the X-ray crystal structures.^25-27^

To investigate the potential junction opening further, the binding to an analogous ‘mismatch’ 3WJ structure where the central cavity has a mismatched T-T base pair (Figure 3 – mm-3WJ) was studied. This structure should promote 3WJ junction cavity expansion and allow the Au pillarplex to have a more comfortable fit, but it will also reduce the underlying stability of the 3WJ structure.^58–61^ This is reflected in the binding of the cylinder (Figure 3 lanes 15-18) where binding to both mm-3WJ and 2-stranded Y-shaped fork structures are now observed even at low loading, indicating that the mismatch has made the junction less attractive to the cylinder. By contrast, the binding of the Au pillarplex to the mismatched-3WJ is very similar to its 3WJ binding with initial mm-3WJ binding observed (and increased hydrodynamic radius) followed by a transition to fork binding at higher loading (Figure 3 lanes 7-10).

The data confirm that the Au pillarplex can bind to and stabilise both 3WJ and Y-fork structures. The binding to 3WJ is stronger than to Y-fork (as expected purely on electrostatic grounds) but both Y-junctions (3WJ or fork) are preferred over binding to single-stranded DNA. The similarity between the Au pillarplex’s observed binding to both 3WJ and mm-3WJ is consistent with the supramolecule expanding the 3WJ cavity (breaking some base-pairing at the junction).

DNA-melting experiments (monitored by the change in UV-absorption^62^) with the 3WJ formed from 14-base sequences as used for the gels, showed no melting curve in the absence of the metallo-supramolecular cations because the 3WJ is unstable at room temperature and single stranded DNA is present throughout. By contrast, a DNA melting curve is observed in the presence of both the pillarplex and the cylinder (Figure S3). The melting temperature (T_m_) is 56.2 (±0.1) °C with the Ni cylinder and 48.8 (±0.5) °C with the Au pillarplex indicating a greater stabilisation of the 3WJ structure when the cylinder is bound which is consistent with the pillarplex binding breaking base-pairs. Using longer 18-base sequences, melting of the free 3WJ is observed (38.5 (±0.7) °C) and a similar greater stabilisation is observed for Ni cylinder (T_m_ = 59.8 (±0.6) °C) than Au pillarplex (T_m_ = 56.5 (±0.6) °C) (Figure S4).

The pillarplex has D_2d_ symmetry with a total of 12 aryl rings arranged in four sets of three, each comprising two imidazoles from one ligand and a pyrazole from the other bridging a pair of coinage metal ions (Figure 1 and S5). Given this tetragonal arrangement of the pi-surfaces and the indication that the pillarplex is a little too wide for the perfect 3WJ, we were intrigued to see whether it might also bind a four-way junction structure (4WJ). The open 4WJ form will have a larger central cavity than the 3WJ.

Analogously to the 3WJ gel studies, a 4WJ was assembled from 4 complementary strands (Figure 2) and exposed to increasing concentrations of Au pillarplex (Figure 5). The 4WJ strands were selected as they have been used by Searcey et al previously to study 4WJ binding of organic molecules using gel electrophoresis.^47^ At room temperature the strands form a mixture of 4WJ and individual single strands, with the 4WJ further stabilised by high concentrations of MgCl_2_ and NaCl. A very small proportion of a higher order structure, is also observed with these cations (Figure 5 lane 2).

**Figure 5.**
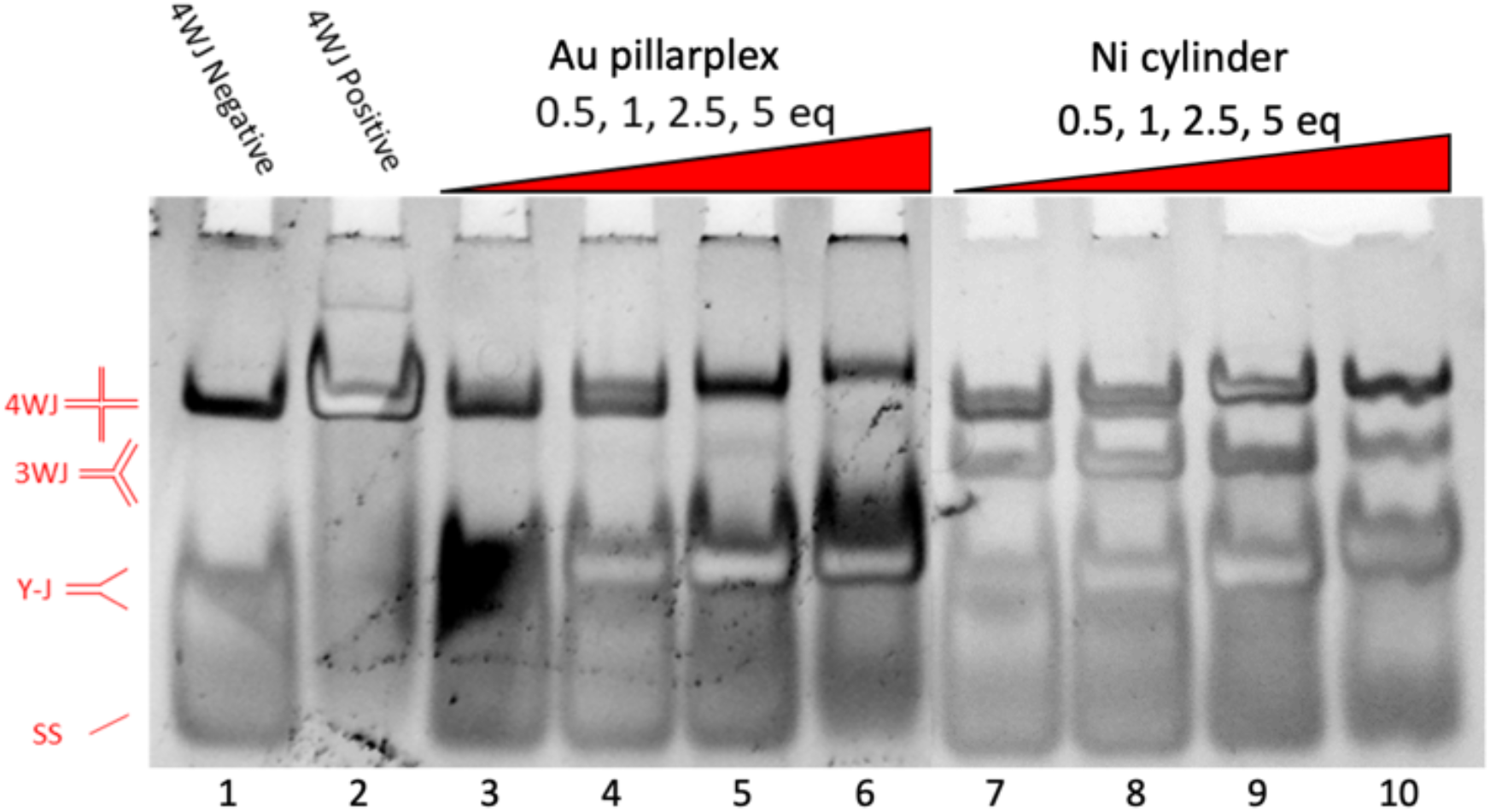
PAGE gels of 4WJ (S1,S2,S3 and S4) incubated with Au pillarplex and Ni cylinder at varying ratios. Controls in lanes 1-2 (lane 1 -strands alone. Lane 2 in presence of cations – 2 mM Mg^2+^ and 450 mM Na^+^).

The Au pillarplex interacts with the 4WJ, causing the appearance of a bound species running slightly slower than the unbound 4WJ. At a 1:1 ratio both are present and the 4WJ band is split into two bands (Figure 5 lane 4). In absence of pillarplex the closed stacked-X junction structure is expected and the band shift is consistent with the bound structure being an open junction and cruciform shaped. As the ratio of Au pillarplex to 4WJ increases, the 4WJ population shifts to favour the bound species band. At high loading the discrete bound band is further retarded, perhaps indicating a second binding can occur, but also reduces in intensity. This decrease in intensity coincides with an increase in intensity of a Y-fork double stranded band, just as seen in the 3WJ gel, indicating again the pillarplex’s preference for fork structures. However, at high loading the 4WJ is still present, (whereas at the same loading, the 3WJ band had disappeared – Figure 3 lane 6) and this suggests that the 4WJ may be a better binding target than 3WJ, likely due to the increased size of the central cavity. A very small proportion of discrete three-strand structure is also observed. Once again, controls with the ligand do not show binding (Figure S6).

We also investigated the binding of the Au pillarplex to the 4WJ strands individually and in all combinations (Figure 6). The pillarplex causes band smearing of the individual strands (S1-3), but for strand S4 we see induction of higher order structures. S4 has the possibility to form a partial duplex containing two 2-base bulges (SI Figures S7-8) and it appears that the pillarplex binds and stabilises this and/or other higher order structures. The other strands do not have the capability to form such a dimer.

**Figure 6.**
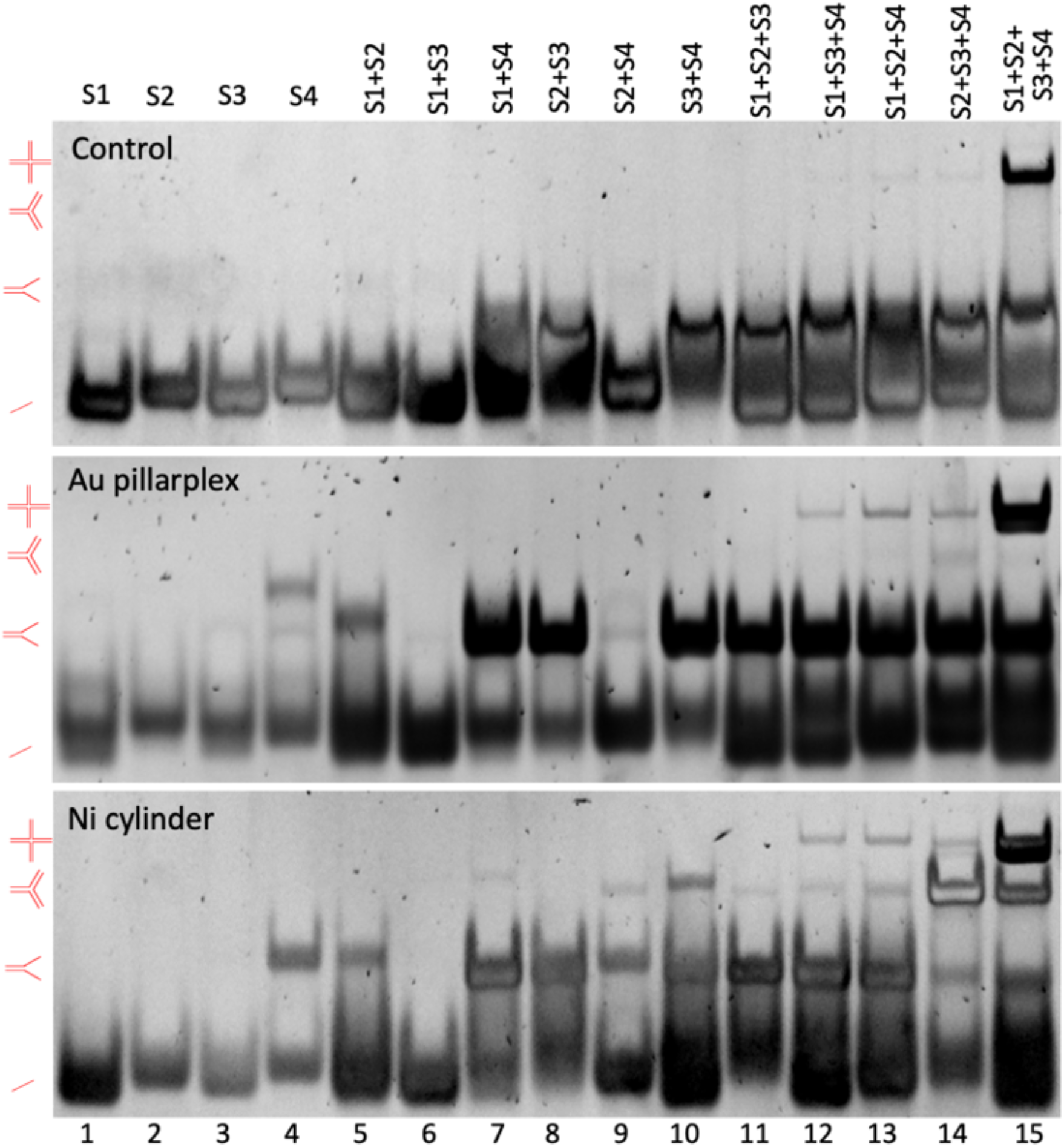
PAGE gels of the individual 4WJ strands (S1,S2,S3 and S4) alone and in combination, alone (top) and incubated with one equivalent of Au pillarplex (middle) and Ni cylinder (bottom).

Where combinations of 2 strands are complementary and can form a Y-fork (i.e. S1S2; S2S3; S3S4; S1S4) the pillarplex stabilises this structure which is consistent with the observed binding to these structures when all 4 strands are present. All four combinations of 3 strands can form a fork-like 3WJ (or bulged 3WJ with partial interactions in one arm -SI Fig. S9) but this seems only stabilised for the S2S3S4 combination and even then in only small amounts. Instead when S4 is present other *tetra-*stranded structures are stabilised (likely 4WJ containing two S4 strands -SI Fig. S10). This again demonstrates the pillarplex preference for 4WJ and duplex forks.

By contrast, when the cylinder [Ni_2_L_3_]^4+^ interacts with the 4WJ structure, the most striking feature is the immediate formation of 3WJs and duplex Y-forks (Figure 5). There is some splitting of the 4WJ band indicating that the cylinder can also bind to the 4WJ (and that bound and unbound species 4WJ are present) but the cylinder preference for Y-shaped structures (especially 3WJs - SI Figure S9) serves to highlight the greater comparative 4WJ preference of the pillarplex. Studies with the individual strands and their combinations (Figure 6) reinforce this observation, with the cylinder also promoting the formation of 3WJ structures in those experiments, including in two strand experiments when S4 is present.

MD simulations of Au pillarplex with a 4WJ DNA model (based on pdb 1XNS) were conducted.^63^ The DNA structure is a crystallographically characterised example in complex with Cre proteins and small peptides (which we removed before introducing pillarplex) and was selected as a starting point for the simulations because its junction cavity is partially open. The starting junction cavity is rectangular in shape and at the junction only two of four potential base pairings are present at the outset (SI Figure S11).

From starting positions with the pillarplex outside, but close to (∼1nm), the junction cavity, the pillarplex was observed to rapidly enter the cavity. The junction then rapidly rearranged to a more open and square form - an open 4WJ structure with the four DNA arms distinct and not stacked together (Figure 7). This is consistent with the small gel band-shift observed. As the cavity re-arranged about the pillarplex, some transient breaking of pairs and stacking of individual bases around the pillarplex was observed as illustrated in Figure 7 (top). However as the simulations proceeded full base pairing at the junction point was recovered and retained (Figure 7 bottom). Importantly, during all the simulations (three at 2 μs each) the pillarplex remained within the central cavity throughout. In simulations in the absence of pillarplex, the free DNA rapidly closed up into the closed Holliday junction form with stacked coaxial arms as would be expected (SI Figure S12). In simulations starting from this closed form, we did not see pillarplex insertion over three 1 μs simulations though this is unsurprising given the timescales of the dynamic opening and closing of a DNA 4WJ. This illustrates the power of sampling different known DNA starting conformations to explore the complex conformational landscapes of nucleic acids even within relatively short timescales afforded by MD simulations, and is similar to simulation approaches we explored previously for RNA binding of cylinders.^55^

**Figure 7.**
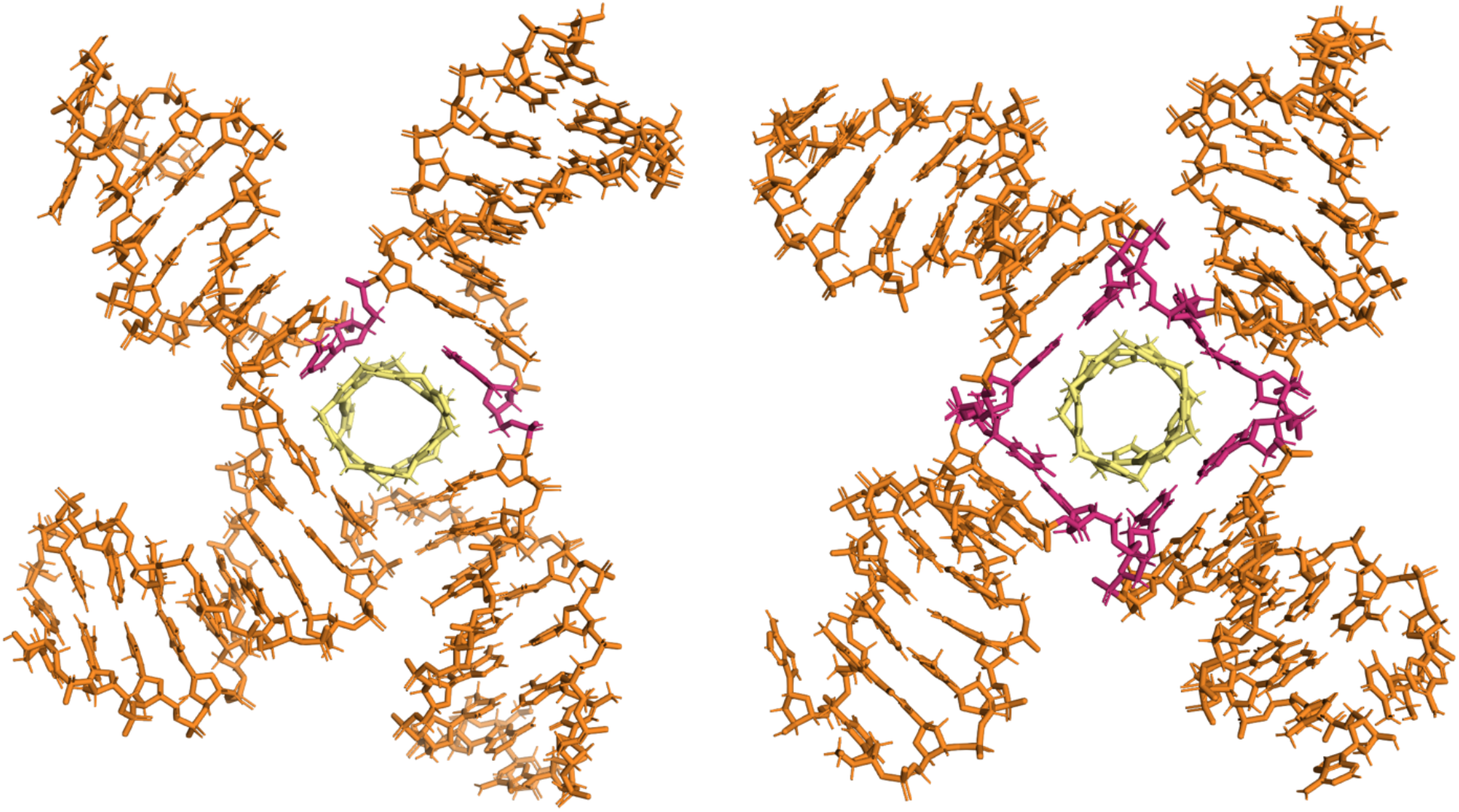
MD simulations of Au pillarplex with a 4WJ DNA. The MD shows Au pillarplex entering and residing in the central cavity of the 4WJ with the cavity closing up around the pillarplex but an open 4WJ structure resulting with the four arms distinct and not stacked together. Following the entry of the pillarplex, much of the base pairing at the junction point is maintained though initially there is transient breaking of pairs and stacking of individual bases around the pillarplex as the cavity flexes to accommodate the pillarplex. This is seen in the left hand image where two bases (highlighted in pink) have become unpaired. As the simulations proceed the pairings reform and an open 4-way junction with fully paired bases contains the pillarplex. This is seen in the right hand image where the 8 bases at the junction are highlighted (pink)

While junctions are an exciting target, the majority of genomic DNA is found in duplex form so we also explored the effects of pillarplex on duplex DNA, starting with a double-stranded DNA (DS21) formed from two complementary strands (Figure 2). In a gel electrophoresis experiment (Figure S13), the Au pillarplex caused a retardation in the duplex band, indicating an interaction, along with some smearing. The cylinder shows a similar (though less marked) effect while the metal-free pillarplex ligand showed no binding. Similar effects were seen with a palindromic 24-mer (Pal24). Likewise, with a longer, biological DNA (pBR-322 plasmid DNA - 4361 base pairs) the linear form showed the same response of a sequential retardation as more Au pillarplex is introduced (Figure S14). DNA precipitation in the well was also observed at high loading, as for the cylinder which is known to coil and condense duplex DNA.^65–67^ The supercoiled, closed circular form of the plasmid DNA, showed unwinding on Au pillarplex binding, with the extent again similar to that of the cylinder.^68,69^

To probe this duplex interaction in more depth we used flow linear dichroism (LD) with calf-thymus (CT) DNA, as a representative genomic DNA. Flow LD allows the orientation of the DNA and the complex to be probed spectroscopically in a Couette cell by comparing the absorption of linearly polarised light parallel and perpendicular to the direction of flow.^65,66,69^ DNA alone shows a strong negative signal at 260 nm due to the DNA bases that become oriented as the DNA helix orients in the flow. The cylinder coils the DNA and this leads to a loss of orientation and a dramatic decrease in the magnitude of the 260 nm signal.^65,66^ The same effect is observed with the Au pillarplex at low loading (Figure 8), however at higher loadings (from 1 pillarplex per 20 base pairs) a second effect is superimposed as a strong positive signal appears. The signal might arise from pillarplex spectroscopy^54c, 70^ (Figures S17-18) or from the DNA bases (or both) but the chromophores are no longer orientated perpendicular to the flow direction but more parallel to the flow orientation, perhaps indicating coiling into some sort of tube conformation. The corresponding circular dichroism spectroscopy shows the B-DNA structure retained at low loading, but the spectra becoming more complex at higher loading which is likely to represent pillarplex induced CD signals overlain on the DNA spectroscopy. The gel and spectroscopic studies show that the pillarplex is condensing duplex DNA, as other polycations do, but may induce an ordered condensed structure.

**Figure 8.**
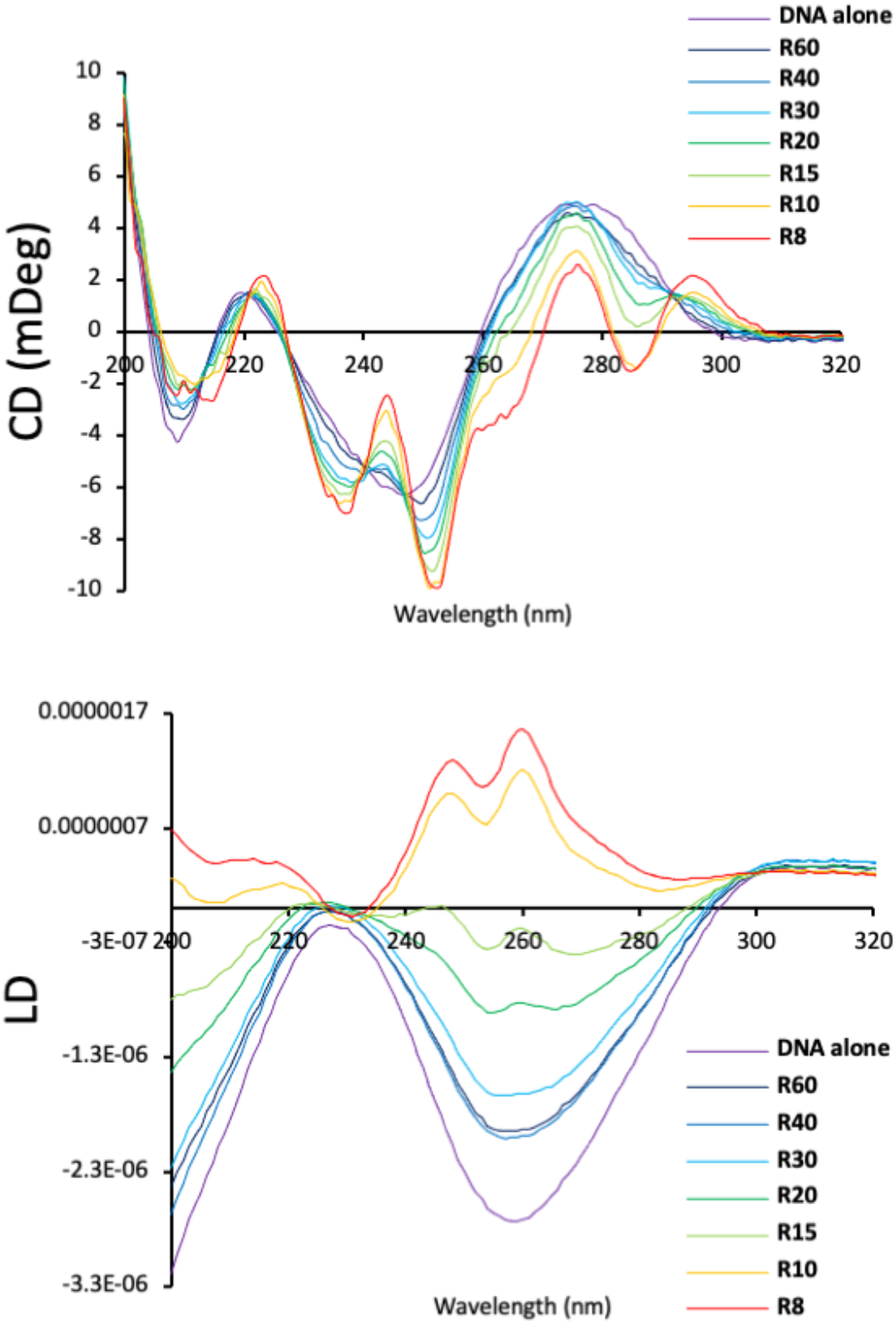
Circular dichroism (top) and flow linear dichroism (bottom) spectroscopic studies of Au pillarplex with CT-DNA. The R value is the ratio of DNA base pairs to complex.

Molecular dynamics simulations of multiple pillarplexes and a 25-base pair double-stranded B-DNA (Figure 9) showed positioning of the pillarplexes along the DNA minor groove, with orientation of the pillarplexes inducing changes to the length of the groove, but did not directly inform on any DNA condensation in the timescales that can be accessed through simulations. However, a second frequent binding mode was observed which involved an opening up of the terminal base pairings at the fraying ends of the double-stranded DNA and insertion of the pillarplex inside the DNA helix. Such a binding mode is reminiscent of both the 3WJ simulations where a base pair is opened, and the 4WJ simulations where a base pair initially breaks to better pack the bases around the pillarplex. It is also a likely model of the binding to Y-fork junction structures that are observed in both the 3WJ and 4WJ gel electrophoresis experiments. It is clear from the experimental data that the pillarplex prefers such junction structures over a conventional duplex and so a potential role in the DNA condensation is an interesting possibility.

**Figure 9.**
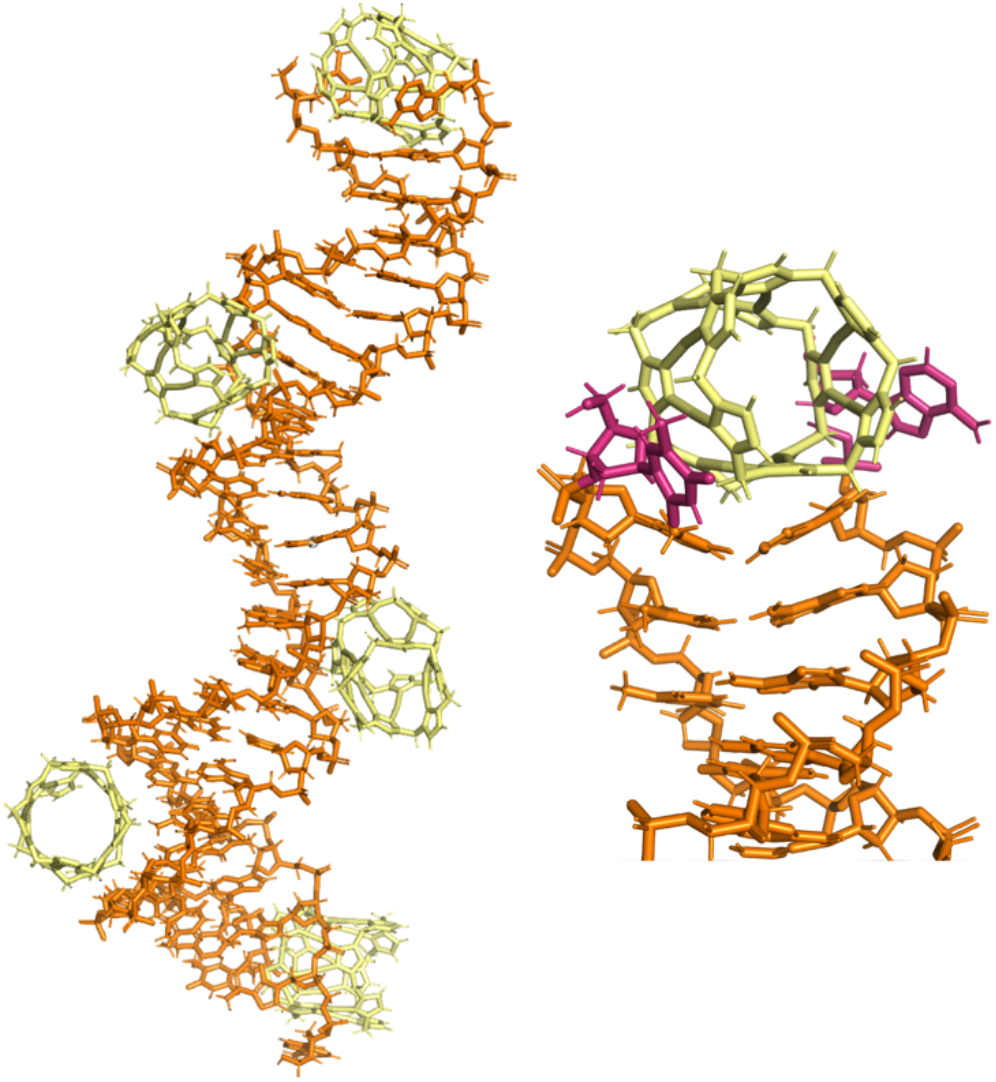
MD simulations of Au pillarplexes with a double-stranded 25 base pair DNA oligonucleotide. Pillarplexes can be seen binding in the minor groove (left) as well as at the fraying ends of the DNA (right – with unpaired bases highlighted in pink).

We have also studied the DNA binding of the corresponding Ag pillarplex structure (SI Figures S2, S6, S13-15). This silver organometallic complex has the same shape and charge as the Au pillarplex, but a lower stability.^70^ Potential complex degradation and a lower water solubility are complicating factors in the experiments, but the DNA binding is similar to that of the Au pillarplex confirming the importance of the structural motif in the DNA binding.

To explore whether the Au pillarplex can enter the cell and access cellular DNA, we explored the uptake and accumulation in A549 lung cells by treating with the compound (6 μM) for 24 h, and quantifying the intracellular metal content by Inductively Coupled Plasma Mass Spectrometry (ICP-MS), and also fractionating to isolate the nucleus and assess its Au content. The initial results confirm that the Au pillarplex can enter the cell with the amount of Au accumulated in the cell extract (ca. 0.08 ng metal/μg protein) very similar to the amount of platinum from cisplatin (used as a control). Approximately 10% of that Au is found in the nuclei after 24 h, which is again comparable with cisplatin.

Antiproliferative activity studies conducted on both Au and Ag pillarplexes against human A549 lung and SKOV-3 ovarian cancer cell lines are reported in SI Table ST1. At 24 h the Ag pillarplex shows some cytotoxic activity (IC_50_ 6-10uM) with a similar activity observed at 72h. By contrast the Au pillarplex is not cytotoxic at 24h (despite having already entered the nucleus) but is moderately cytotoxic at 72h (IC_50_ 11-12uM) at levels comparable to those of cisplatin.

## CONCLUSION

We have shown herein that the pillarplex binds to open cruciform-style 4WJs which is exciting, both because these are believed to be the biologically active form of the Holliday junction,^7^ and because this now provides a roadmap for targeting higher order junction structures using this metallo-supramolecular approach.

The increased girth of the pillarplex compared to the previously studied cylinders brings new properties to its binding: Although the pillarplex does bind 3WJ, in doing so it disrupts the hydrogen bonding between bases and opens up that junction structure, and at higher loading it breaks apart the 3WJs into Y-shaped forks. Indeed, the contrast with the cylinder is striking, with the cylinder rearranging the 4WJ into its preferred 3WJ targets (or Y-forks) whereas the pillarplex does not transform 4WJs into 3WJ structures.

That the Au pillarplex is an effective junction and fork binder, indicates the value of developing cationic agents which display polyaryl surfaces as a junction-binding design strategy. Featuring an organometallic scaffold, and a pore (that can be used for rotaxane formation)^54^ it further expands the chemical toolbox available to construct nucleic acid junction binders.

The pillarplex is a little small to completely fill an open 4WJ cavity and there is scope for further elaboration to create slightly larger agents to target and reside in this cavity. This minor size-mismatch helps to explain the competition between 4WJ-binding and Y-shaped fork binding observed at high pillarplex loading.

The 24h timepoint where the Au pillarplex has entered the nucleus but not yet caused cytotoxicity offers the opportunity for future detailed study of the *in cellulo* effects of Holliday junction binding. Alongside their biological roles, DNA junctions – and especially 4WJs – are also and widely used in nucleic acid nanostructure construction. The ability of the pillarplex to not only form and stabilise DNA junctions, but also modulate and switch them is particularly intriguing in this context. We are actively exploring these aspects.

## SUPPORTING INFORMATION

The Supporting Information includes: Experimental details, Supplementary Table ST1 & Supplementary Figures S1-S19 (PDF)

## ACKNOWLEDGMENTS

This work was funded by the EPSRC Physical Sciences for Health Centre (EP/L016346/1) BBSRC MIBTP (BB/M01116X/1), DFG-SPP 1928, the Swiss National Science Foundation (200020_165868; 200020_192153), University of Zurich and the University of Birmingham. Simulations used the Bluebear and Castles HPC facility (U. Birmingham).^71^ We thank the Institute of Advanced Study, TU Munich and the Institute of Advanced Studies, U. Birmingham for network meetings to establish the collaborations. AP was supported by a Vanguard visiting fellowship from the University of Birmingham Institute of Advanced Studies. We thank Dr Zenghui Wang and Dr Ilaria Przytula-Mally (Zurich) for support in nucleic acid NMR studies and crystallisation attempts. We thank Dr Anton Vladyka and Prof Tim Albrecht (Birmingham) and Dr Stefano Leoni (Cardiff) for helpful conversations and studies.

## CONTRIBUTIONS

MJH and AP conceived, supervised and directed the project. JSC undertook the gel and spectroscopic experiments, and designed experiments with MJH. HW undertook gel and UV melting experiments and CC undertook gel experiments. DFB supervised radio-labelled gels. LM and SD undertook the MD simulations. RKOS, SJ and RE investigated oligonucleotide NMR and crystallization studies. AP, AAH, PJA and SG prepared the pillarplexes and ligand, and JSC prepared the cylinders. AC and BA contributed the antiproliferative activity and cellular accumulation studies. MJH and JSC drafted the manuscript which all authors commented on.

