## Supporting Information for "Organometallic Pillarplexes that bind DNA 4-way Holliday Junctions and Forks"

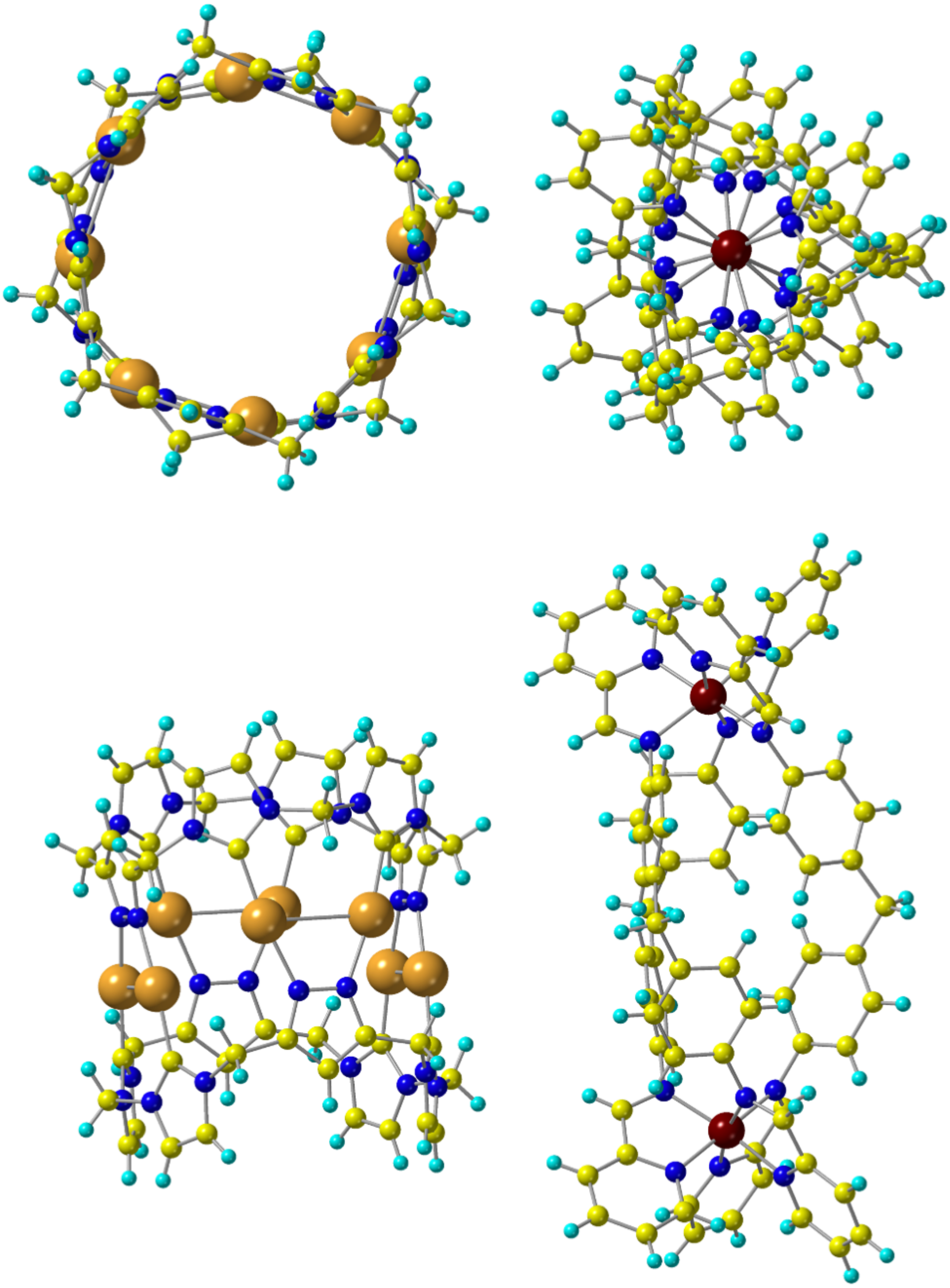

**Figure S1a** Comparative views of the pillarplex (left) and cylinder (right) from the top and side.

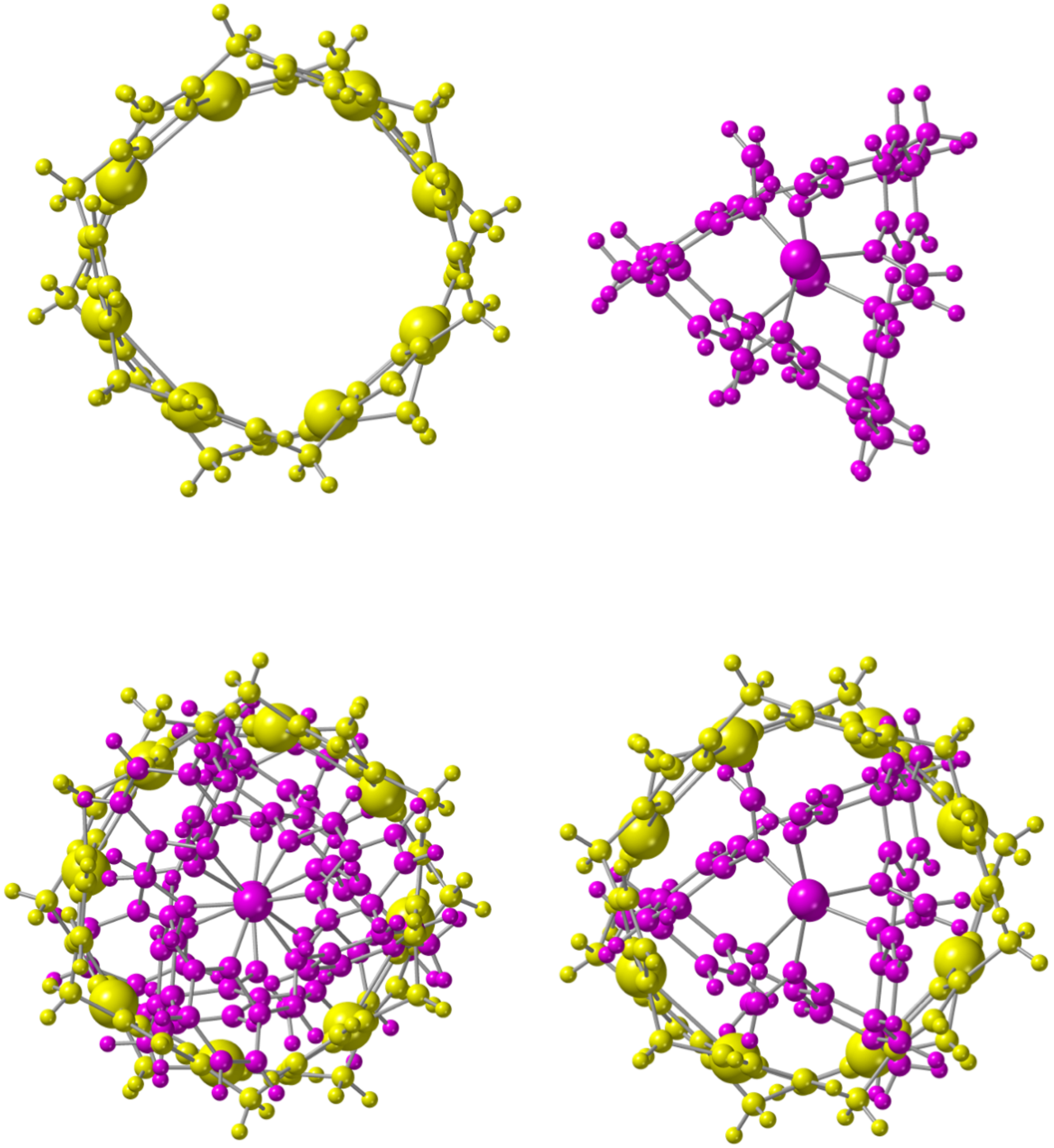

**Figure S1b** Comparative views of the pillarplex (top left) and the central core of the cylinder which binds the 3WJ cavity ie with the pyridyls cut away (top right). Overlay of the full cylinder structure with the pillarplex (left) and just the central core of the cylinder with the pillarplex (right).

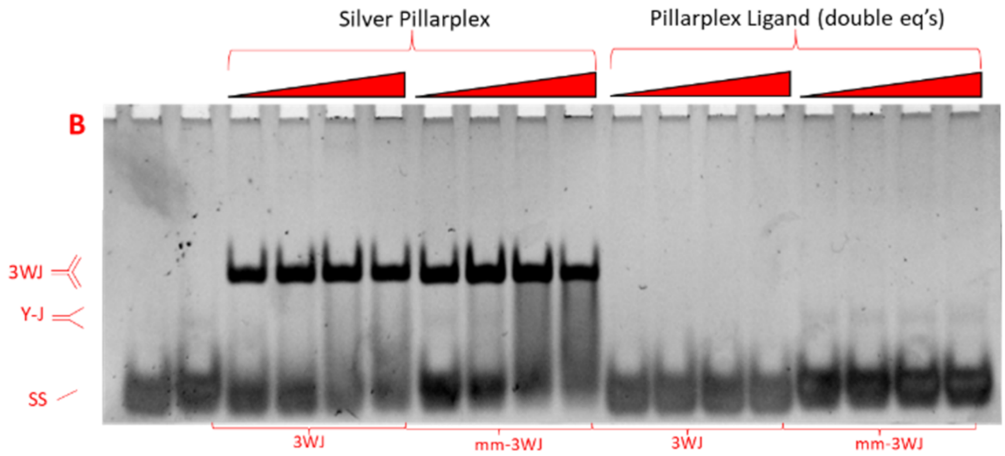

**Figure S2:** Shows a PAGE gel of 3WJ and mm-3WJ DNA strands incubated with Ag pillarplex and pillarplex ligand at varying ratios. The gel is analogous to main paper Figure 3 (Au pillarplex and Ni cylinder). Gel lanes read from left to right. Controls in lanes 1 (3WJ strands alone) and 2 (mm-3WJ strands alone). 3WJ is mixed with Ag pillarplex or ligand at 0.5,1, 2 and 4 eq (lanes 3-6 and 11-14), and mm-3WJ mixed with Ag pillarplex or ligand at 0.5,1,2,4 eq (lanes 7-10 and 15-18).

The Ag pillarplex stabilises the 3WJ and mm-3WJ structures. In both cases a shifted 3WJ band is observed which is very similar to that observed with Au pillarplex. At high loading some smearing below the 3WJ is observed, however (in contrast to Au pillarplex) we do not see a pronounced Y-shaped band at higher loading. The Ag pillarplex has poorer solution stability than the Au pillarplex, and there is potential for release of silver cations (from pillarplexes not bound in the 3WJ cavity) which may then bind to the DNA.

The free polyaryl ligand from which the pillarplexes are constructed shows no binding to DNA.

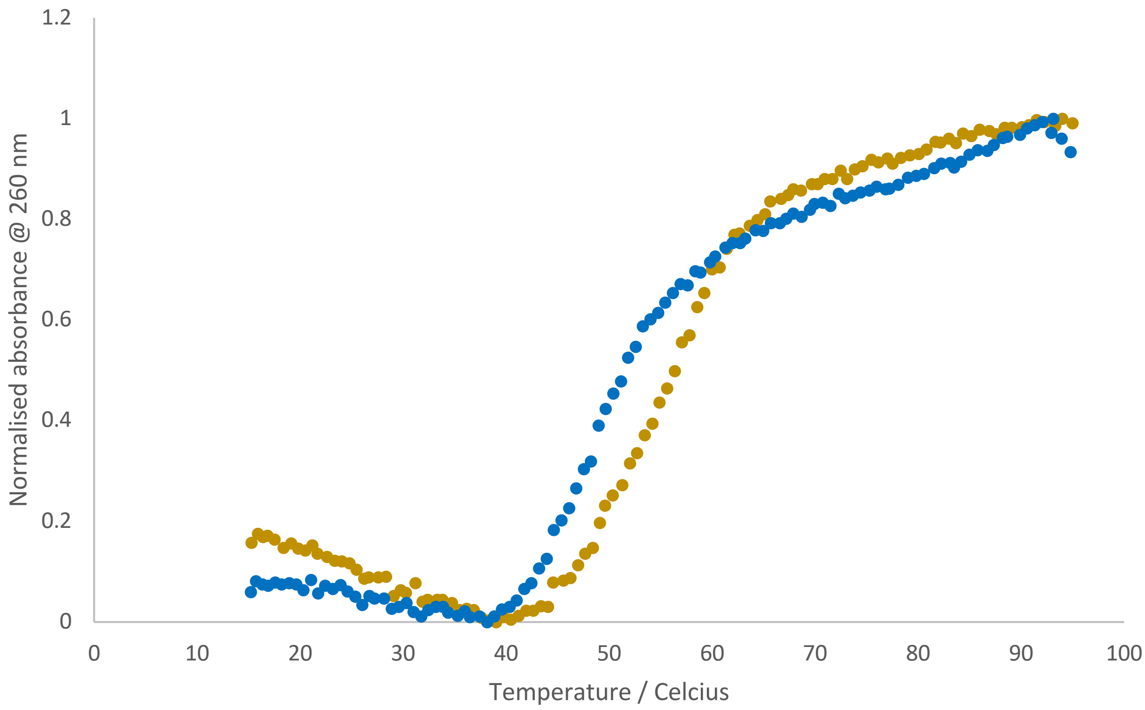

**Figure S3:** Representative data for the 3WJ UV-Vis melting experiments (14 bases per strand) with Ni cylinder (yellow) and with Au pillarplex (blue).

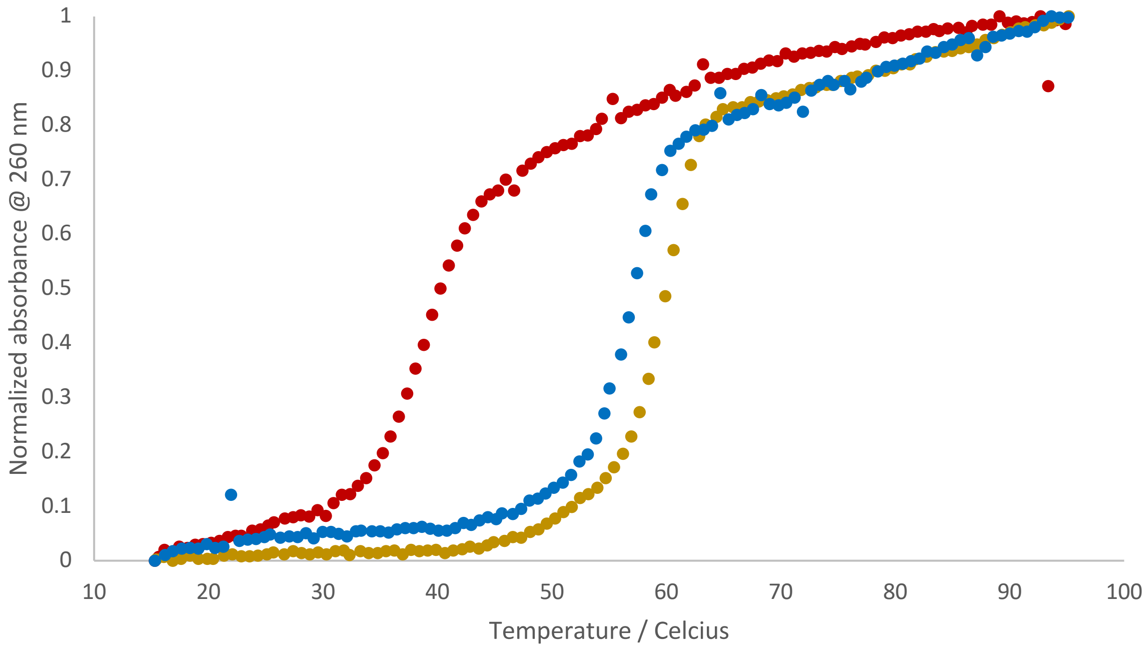

**Figure S4:** Representative data for 3WJ18 UV-Vis melting experiments alone (red) with Ni cylinder (yellow) and with Au pillarplex (blue)

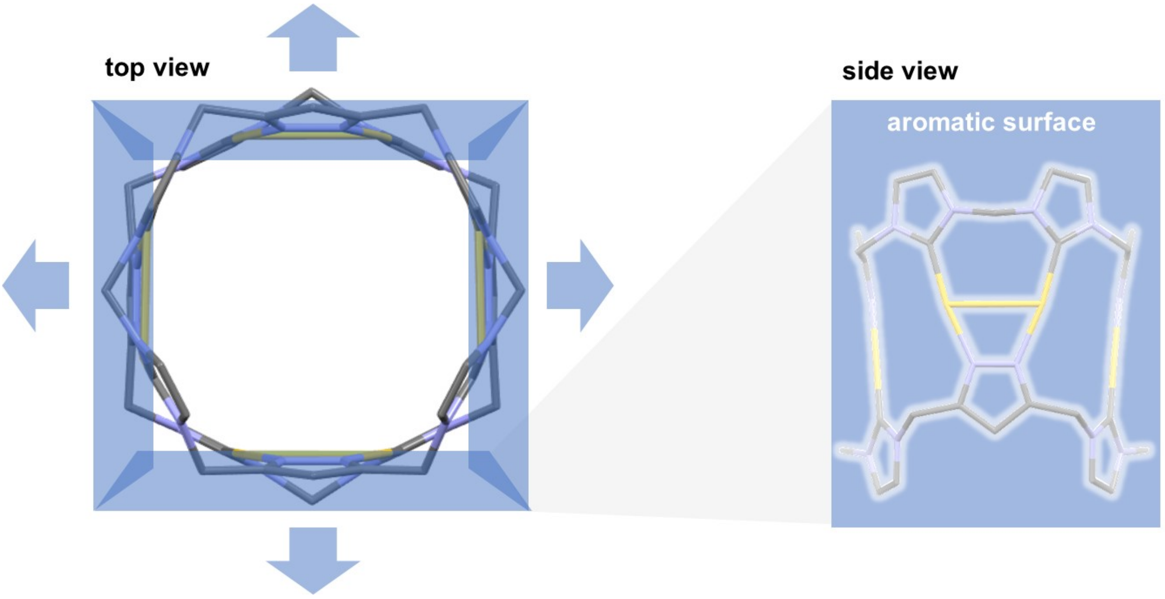

**Figure S5** Views of the pillarplex emphasising the tetragonal arrangement of surfaces within the structure

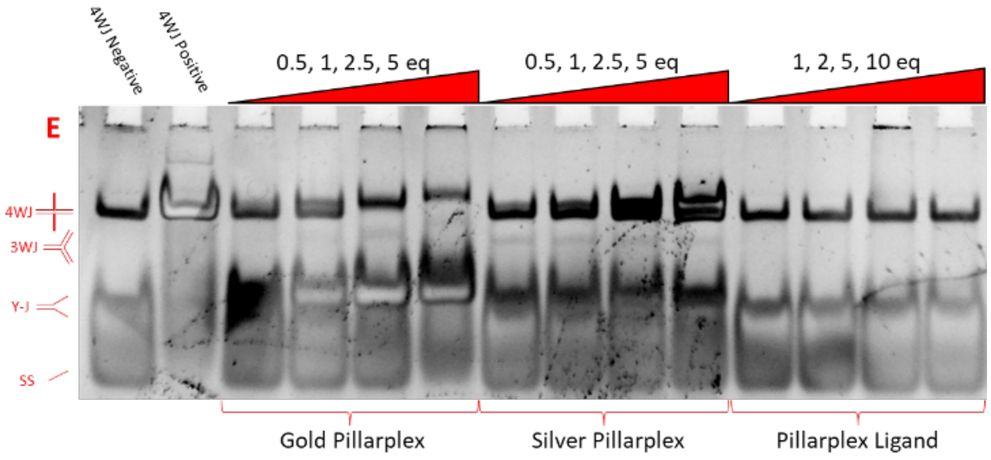

**Figure S6** Shows a PAGE gel, of a 4WJ incubated with Au and Ag pillarplexes and the free pillarplex ligand at varying ratios. Lanes 1-6 (left to right) are also shown in the main paper Figure 5. S1,S2,S3 and S4 are present in all lanes, with lanes 1 and 2 representing controls (Lane 1 - strands alone. Lane 2 - in presence of cations 2mM $Mg^{2+}$ and 450mM $Na^{+}$). Complexes or ligand are added in at 0.5,1,2.5 and 5 eq. ratios (compound: 4WJ). Lanes 2-6 Au pillarplex, lanes 7-10 Ag pillarplex and lanes 11-14 pillarplex ligand.

The Ag pillarplex (as the Au pillarplex) binds the 4WJ (for which the band broadens and is slightly shifted), with a faint band also observed for the discrete three-strand structure and a larger band for the Y-fork duplex DNA. It is interesting that the 4WJ band experiences less shift at high loading than with Au pillarplex, though this may reflect a lower pillarplex concentration given the lower stability of the silver complex. Once again controls with the ligand do not indicate free-ligand binding.

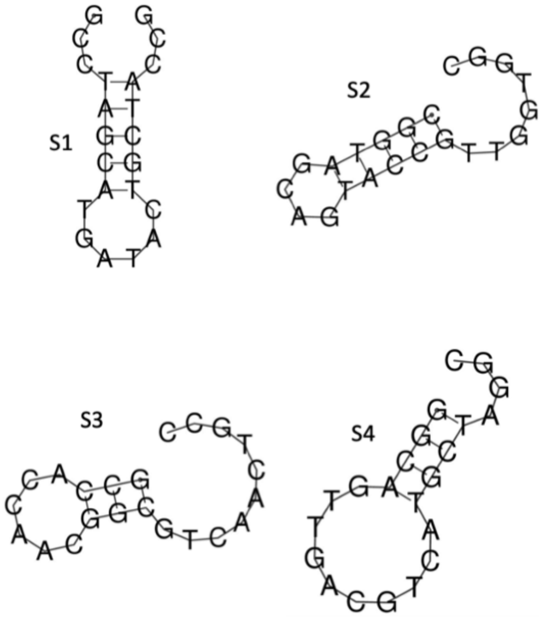

**Figure S7:** Example of how the individual single strand oligos from the 4WJ might fold. Produced using the nucleic acid Fold structure prediction software from the Matthews Lab at U. Rochester available at:
https://rna.urmc.rochester.edu/RNAstructureWeb/Servers/Predict1/Predict1.html
Although the four oligos are the same length, some small differences in mobility are observed in the gel in main paper Figure 6 likely indicating different levels of folding.

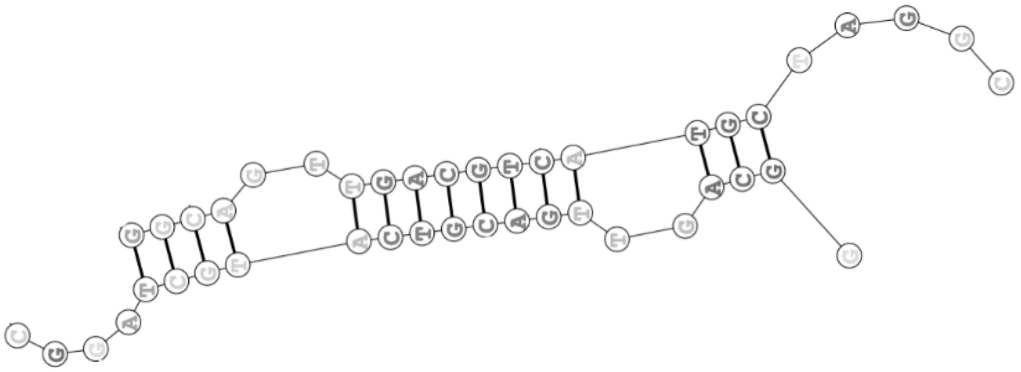

**Figure S8:** Example of how two S4 oligos might interact to form a dimer. The gel in main paper Figure X indicates that S4 produces dimer in absence of pillarplex. Produced using the nucleic acid Fold structure prediction software from the Matthews Lab at U. Rochester available at:
https://rna.urmc.rochester.edu/RNAstructureWeb/Servers/Predict1/Predict1.html
The prediction used a sequence of two S4 strands connected by an XXXXX spacer which was then removed from this graphical representation.

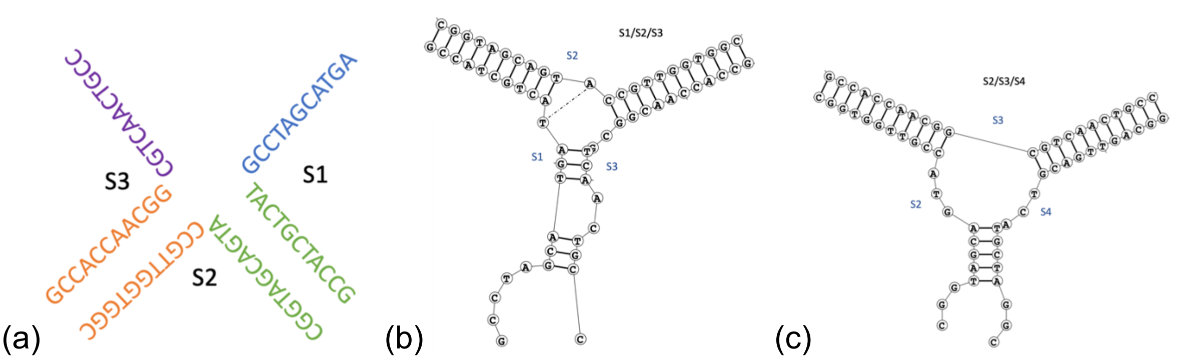

**Figure S9:** Examples of how three oligo sequences might interact to form a 3WJ. The 3-strand bands observed with cylinder may represent formation of an open forked version of the 4WJ (S6a) but more likely the ss arms of that structure will interact (S6b) in this example potentially leading to a 3WJ with a 2-base bulge. This is especially true when cylinder is present to bind in the cavity and stabilise the 3WJ form. Previous work has confirmed the ability of cylinders to bind to 3WJ containing bulges at the junction point as well as perfect 3WJs (Malina, J.; Hannon, M.J.; Brabec, V. Recognition of DNA Three-Way Junctions by Metallosupramolecular Cylinders: Gel Electrophoresis Studies. *Chem. Eur. J.* **2007**, *14*, 3871-3877) Structures produced using the nucleic acid Fold structure prediction software from the Matthews Lab at U. Rochester available at:
https://rna.urmc.rochester.edu/RNAstructureWeb/Servers/Predict1/Predict1.html
The prediction used the sequence of the three strands connected by XXXXX spacers and combined into a single strand, with the XXXXX spacers removed from this graphical representation.

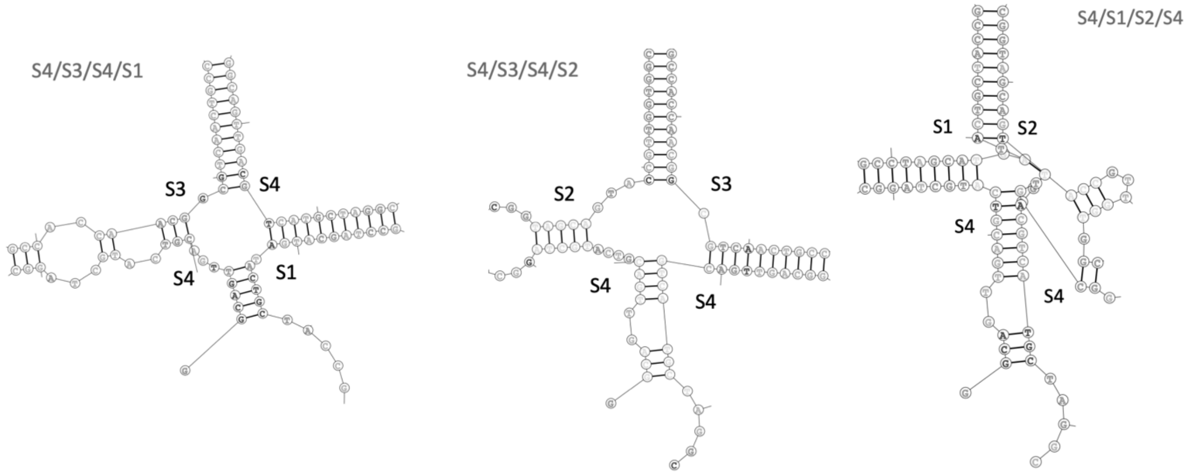

**Figure S10:** Example of how three oligo sequences might interact to form a tetramer. The gel in main paper Figure 6 indicates that combinations including S4 can produce small amounts of a tetramer in absence of the fourth sequence strand, with this seen very weakly in absence of pillarplex, and more strongly in presence of pillarplex or cylinder. Produced using the nucleic acid Fold structure prediction software from the Matthews Lab at U. Rochester available at:
https://rna.urmc.rochester.edu/RNAstructureWeb/Servers/Predict1/Predict1.html
The prediction used a sequence of two S4 strands and two others with each sequence connected together into a single strand by an XXXXX spacer which was then removed from this graphical representation.

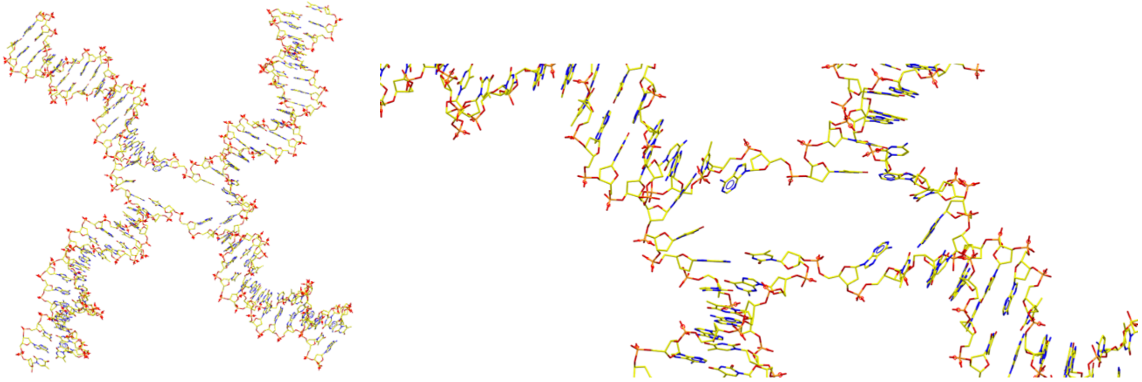

**Figure S11:** Starting position of the DNA 4WJ as extracted from pdb 1XNS (crystal structure) and used in simulations, illustrating the partially open cavity. The second image is a close-up (slightly rotated) to better illustrate the partially open cavity.

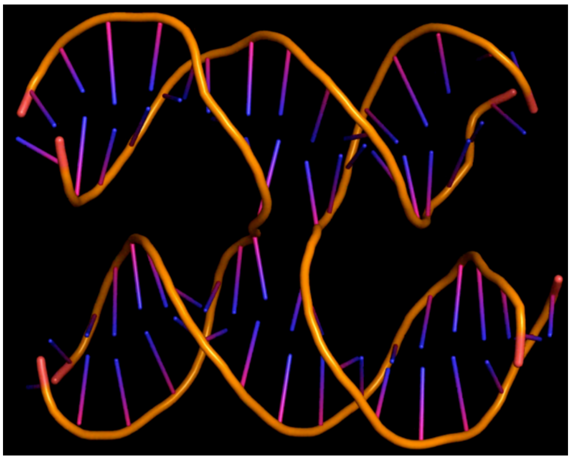

**Figure S12:** Example of the closing of the DNA 4WJ into the closed form in simulations of the free DNA, in absence of pillarplex.

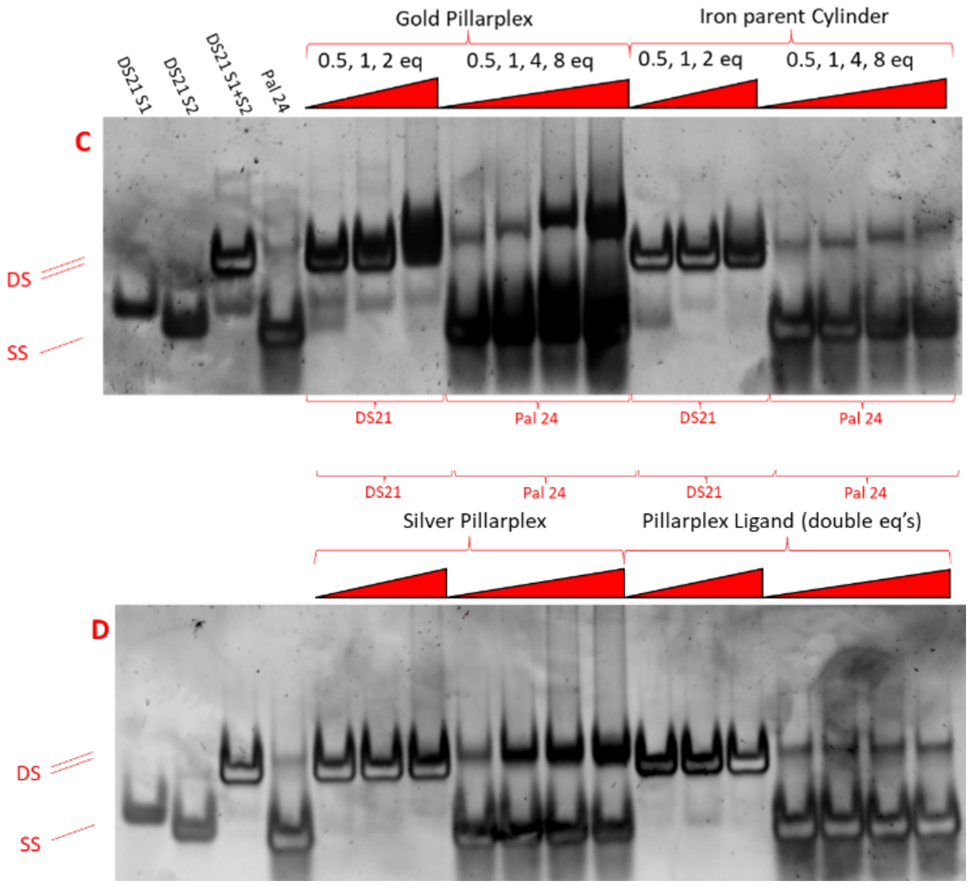

**Figure S13.** PAGE gels of DS21 and Pal24 incubated with gold Pillarplex, iron Cylinder, silver Pillarplex and Pillarplex ligand at varying ratios. Controls in lanes 1-4, then DS21 mixed with a complex at 0.5,1 and 2 eq (lanes 5-7 and 12-14), and Pal24 mixed with a complex at 0.5,1,2,4 eq (lanes 8-11 and 15-18). The silver pillarplex promotes the formation of duplex DNA for Pal24 but its effects are less striking than those of the Au pillarplex. The pillarplex ligand shows no evidence of binding to the DNA.

.

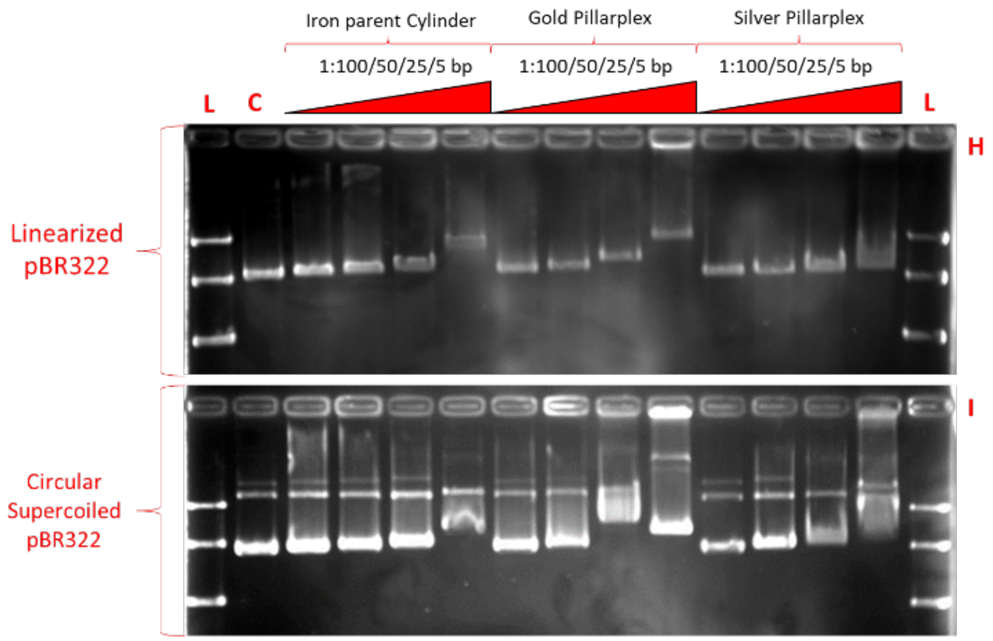

**Figure S14.** Agarose Gel Electrophoresis Studies of Circular supercoiled (bottom) and linearized (top) pBR-322 plasmid DNA with varying ratios of Fe cylinder, Au pillarplex and Ag pillarplex. Lanes 1 and 15 are a DNA Ladder (L), and lane 2 the plasmid DNA alone as control. The Ag pillarplex shows similar behavior to the gold pillarplex but at higher loading values, consistent with some degradation of Ag pillarplex in the experiment leading to lower values of pillarplex in solution.

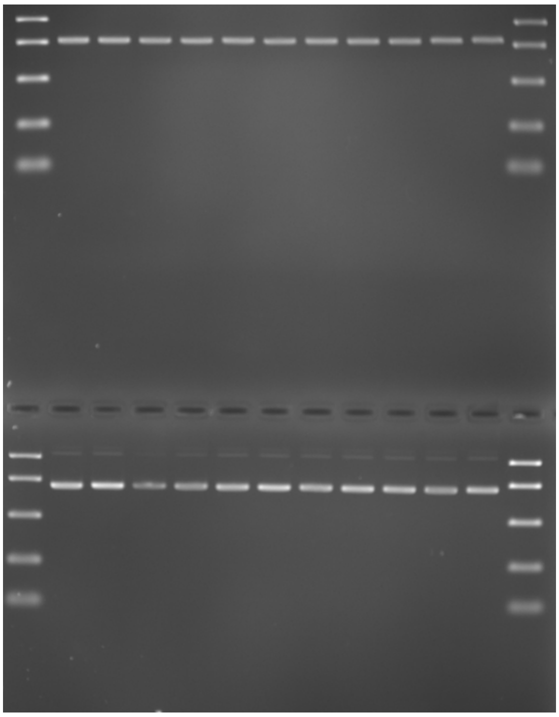

**Figure S15:** Agarose gel electrophoresis gel of pillarplex ligand with PBR-322 linearized plasmid DNA (top) and circular plasmid DNA (bottom). 30:1, 20:1, 15:1, 10:1, 7.5:1, 5:1, 4:1, 3:1, 2:1, 1:1 DNA base pairs to ligand. Ratios are 2x the concentration of pillarplexes in Figure S14, to account for the presence of 2 ligands in each Pillarplex.   No evidence of binding is observed.

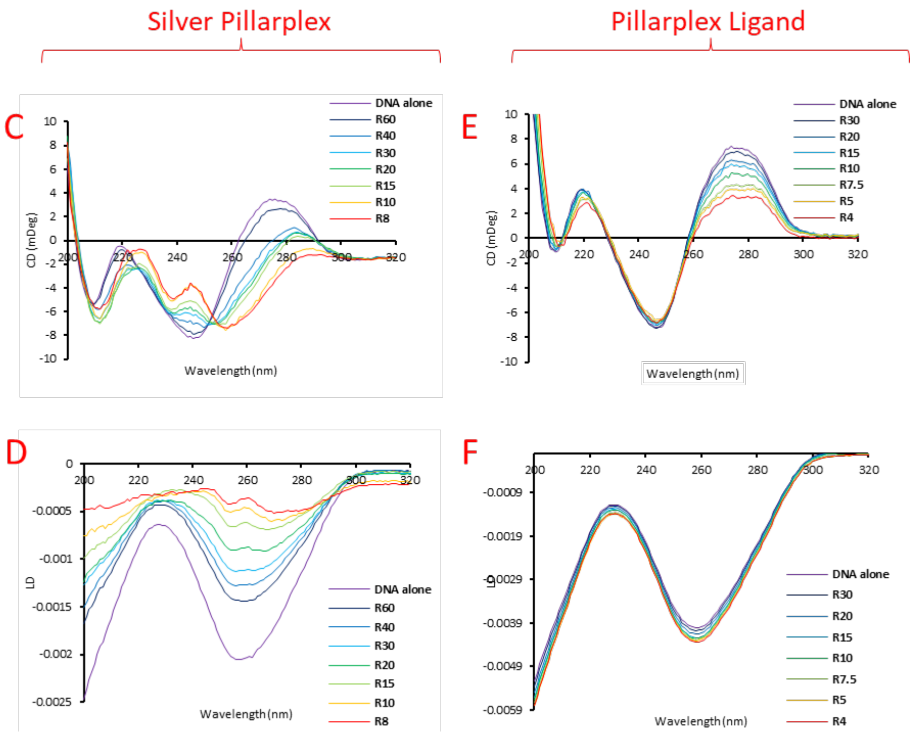

**Figure S16.** Circular dichroism (top) and flow Linear Dichroism (bottom) spectroscopic studies of the of Ag pillarplex and pillarplex ligand with CT-DNA. The R value is the ratio of DNA base pairs too complex. Spectra are acquired by titration of a solution (containing complex of interest) and 2x DNA/buffer solution to an initial solution of 100 μM CT-DNA (bp) with spectra being recorded after each titration addition. The CD and LD spectra of the Ag pillarplex bear similarities to those of the Au pillarplex, though the effects are less striking. In particular in the LD, for Ag pillarplex, the signals do not go positive within this concentration range though if the Ag pillarplex is degraded in the solution the real concentrations may be lower.

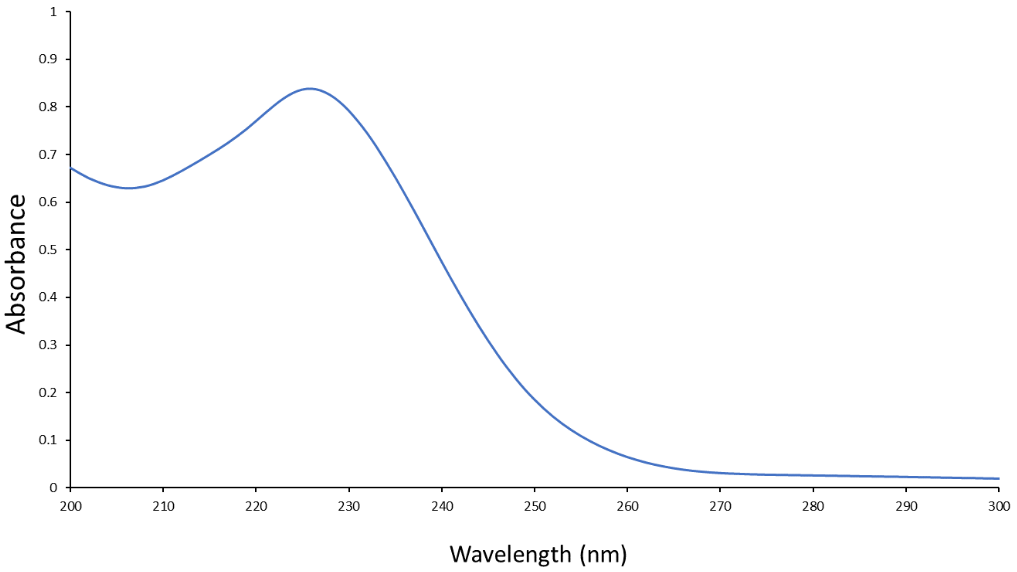

**Figure S17** Ag pillarplex absorbance spectrum at 10uM concentration in water.

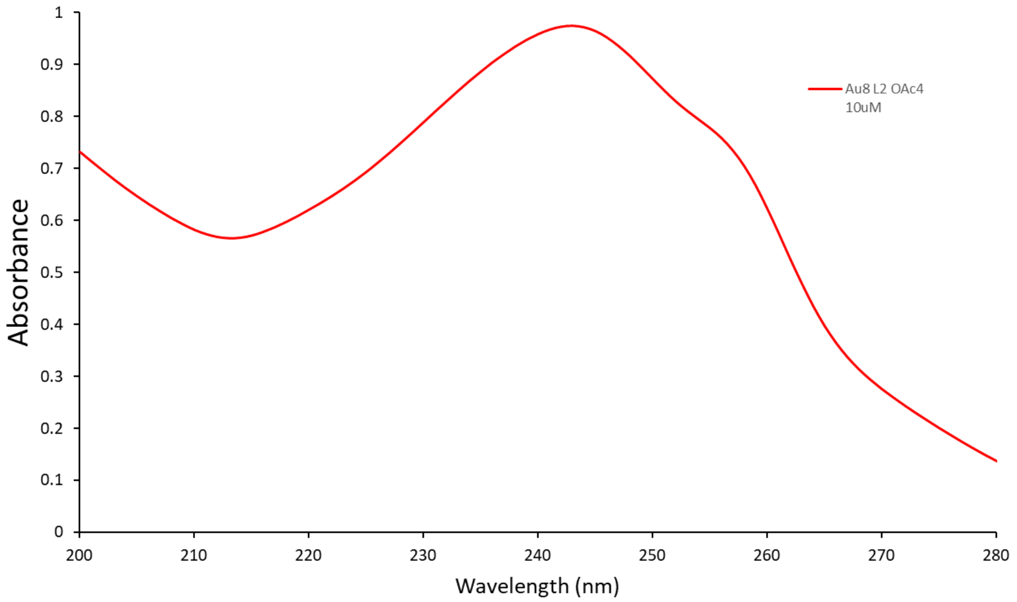

**Figure S18:** Au pillarplex absorbance spectrum at 10uM concentration in water.

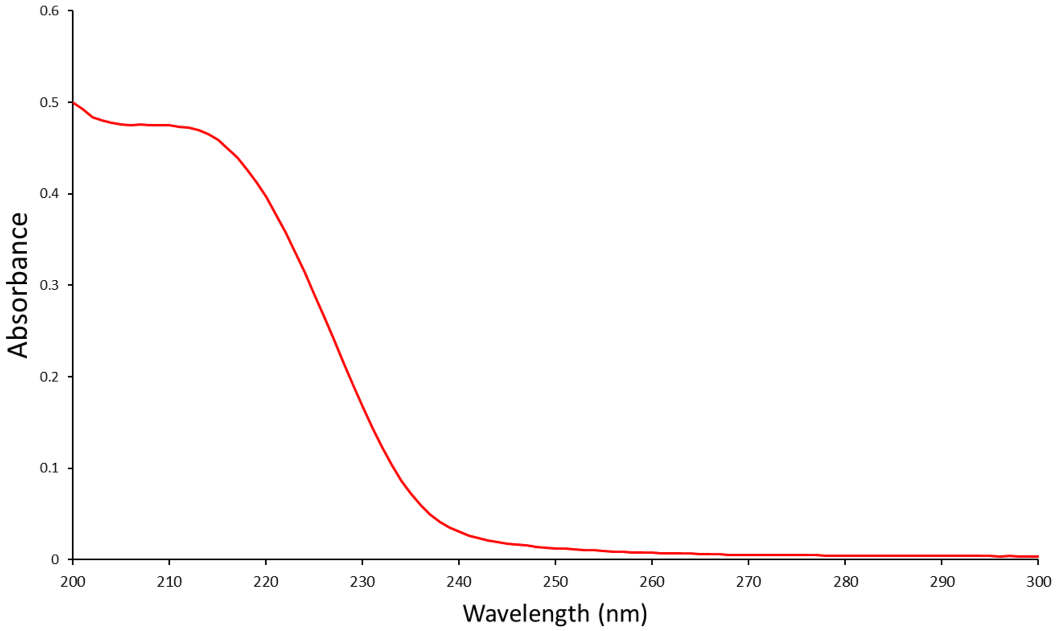

**Figure S19:** Pillarplex ligand absorbance spectrum at 10uM concentration in water.

**Table ST1: Cell cytotoxicity results**

| | Mean $IC_{50}$ ± SD (μM) | | | |
|---|---|---|---|---|
| | A549<br>Human epithelial lung carcinoma | | SKOV-3<br>Human ovarian cancer | |
| | *24h* | *72h* | *24h* | *72h* |
| Au pillarplex | 41.3 ± 4.1 | 11.3 ± 1.8 | >50 | 12.4 ± 1.9 |
| Ag pillarplex | 5.7 ± 2.0 | 6.95 ± 2.85 | 9.7 ± 2.1 | 5.45 ± 0.75 |
| cisplatin | >100 | 11.8 ± 1.5 | >100 | 15.4 ± 2.2 |

### Experimental Details

**Biophysics experimental**: Milli-Q water (18.2 MΩ) was used throughout all biophysical work. For Circular and Linear dichroism, DNA samples were made up from Calf thymus DNA sodium salt (Sigma Aldrich) by dissolving in milli-Q water(18.2 MΩ) and washed using a 10 kDa MWCO centrifuge tube (Sartorius, Vivaspin, 10 ml). The solution was then quantified by UV-Vis spectroscopy (Cary 5000 NIR spectrometer) by ε (260 nm) = 13,200 $mol^{-1}$ $dm^3$ $cm^{-1}$ to give a concentration in DNA base pairs. This stock solution was kept frozen with fresh aliquots being taken out for each experiment. Fresh buffer was made up before each experiment (see experimental for specifics). Complexes were dissolved in Milli-Q water (18.2 MΩ) only with fresh solutions being used for each batch of experiments.

**Circular dichroism (CD**): spectra were recorded on a Chirascan+ spectrometer (Applied Photophysics limited). The samples were scanned in a 1 cm path length cuvette between 800 and 200 nm with 3 repeats at 1 nm step size and 0.5s dwell time per point. Titrations were carried out at a constant concentration of CT-DNA, sodium chloride (10 mM) and sodium cacodylate buffer (1 mM, pH 7.3) by adding compensating solutions of 2x DNA/Buffer of equal volume to the titre of the complex solution. The concentration of complex in the cuvette was increased step wise by adding set volumes of a stock complex solution. The R value refers to the ratio of DNA base pairs to complex, i.e. R60 = 60bp for every 1 complex, R4 = 4bp per complex. CT-DNA concentration was 100 µM in DNA base pairs.

**Flow Linear Dichroism (LD):** was carried out on the same Chirascan+ spectrometer (Applied Photophysics limited) using the LD accessory (Applied Photophysics limited). The LD cell has an angular gap of 0.25 mm giving an overall path length of 0.5 mm. Samples volumes began at 150 µl and stopped at 250 µl. The cell was rotated at 40 revolutions per second to optimise the DNA signal. The titration series was carried out the same as in the CD studies, with a 3-minute incubation time at a lower revolution speed of 4 revolutions per second. CT-DNA concentration was 100 µM in DNA base pairs.

**Polyacrylamide gel electrophoresis (PAGE**): Studies were performed using following oligonucleotide sequences (from Eurofins Germany) which were purified by reverse-phase HPLC.

**DNA melting:** The stability of the DNA three-way junction (3WJ18 and 3WJ) with different junction binding agents was monitored by measuring the absorbance at 260 nm (bandwidth, 1nm; average time 4 s; heating rate, 1.0 ˚Cmin-1; measurement interval, 0.5 ˚C) with increasing temperature. A 1 cm path length, masked, quartz cuvette, and a peltier-temperature controller were used in a Cary 5000 UV-Vis-NIR spectrophotometer. The 3WJ18 structure was composed of 3 separate oligonucleotides, the solutions measured contained 1 μM of each oligo and 1 μM of metal complex and were made up in Sodium Cacodylate buffer (10 mM, pH 7.4) and NaCl (100 mM). The 3WJ melting experiments were carried out in the same manner but with 2 μM each oligo and 2 μM metal complex. The melting temperature (Tm) was calculated using the thermal heating program's built-in smoothing function and first derivative calculation. Each condition was made in triplicate and the $T_m$ reported is the average of three runs.

**DS21**:
S1 = 5' - CCTTCACGCGAACGTAATCCT - 3'
S2 = 5'- AGGATTACGTTCGCGTGAAGG - 3'

**PAL24**:
S1 = 5'-CTTGAGCTTGAGCTCAAGCTCAAG-3'

**b-3WJ:**
S1 = 5'-CGGAACGGCACTCG-3'
S2 = 5'- CGAGTGCTGCGTGG-3'
S3 = 5'-CCACGCTCGTTCCG-3'.

**3WJ**:
S1 = 5'-CGGAACGGCACTCG-3'
S2 = 5'- CGAGTGCAGCGTGG-3'
S3 = 5'-CCACGCTCGTTCCG-3'

**HJ/4WJ**:
S1 = 5'-GCCTAGCATGATACTGCTACCG-3'
S2 = 5'-CGGTAGCAGTACCGTTGGTGGC-3'
S3 = 5'-GCCACCAACGGCGTCAACTGCC-3'
S4 = 5'-GGCAGTTGACGTCATGCTAGGC-3'

**3WJ18**
S1 = GTGGCGAGAGCGACGATC
S2 = GATCGTCGCAGAGTTGAC
S3 = GTCAACTCTTCTCGCCAC

**Fluorescent PAGE experiments**: PAGE Polyacrylamide gels were prepared by mixing 25 ml of 29:1 acrylamide/bis-acrylamide (National Diagnostic, protogel) with 5 mL of 10x Tris-Boric acid buffer (890 mM each, pH 8.3, or pH 7.05 adjusted with HCl) and 20 ml of Milli-Q water. To this 400 μL of a 10% w/v ammonium persulfate/water solution and 40 μL of TEMED were added to initialise polymerisation. This is then immediately poured between 2 glass plates and a comb inserted at the top this is then left to set for 1 hr. Samples were made up to 10 μl containing 1 μM of each strand 1xTBN buffer (89 mM Tris, 89 mM Boric acid, 10 mM

NaCl, either pH 8.3 or 7.05) and the indicated ratio of Complex or competitor. DNA, water, and buffer were mixed in solution before addition of the stated ratios of complex and additional components. Samples were then incubated at 37 degrees Celsius for 1 hr. 5 µl of 30% w/v glycerol was then added to each sample and the sample was then centrifuged, mixed, and pipetted into the wells on the gel. The Gel was run at 140 V for 2 hours in 1xTB buffer. The gel was then removed from the plates and stained using SYBR™ Gold Nucleic Acid Gel Stain (Thermofisher scientifc) in 1xTB buffer for 45 minutes before imaging on a bio-rad ChemiDoc fluorescent imager with 305 nm excitation. DNA structures were at 1 µM in the samples with equal concentrations of each strand needed for their respective structures, complex concentrations are indicated in ratios to this.

**3WJ Radiolabelled PAGE**: To confirm that fluorescent gel results were not affected by any potential fluorescence quenching by metal complexes, 3WJ gels were also repeated using radiolabelled DNA. The results confirmed that SYBR gold staining was sutiable for studying the DNA binding of the Ni cylinder and Au pillarplex in gels. 1.2 µL of S1 was radiolabelled using 1 µL of adenosine triphosphate γ 32P (Perkin Elmer) at the 5' end using 2 µL of T4 bacteriophage polynucleotide kinase (New England Biolabs) by incubating them at 37 degrees Celsius for 1 hr in 2 µl of 10xPNK buffer (New England Biolabs) made up to 20 µL with nuclease free water (ThermoFisher Scientific). This solution was heated to 80 °C for 3 minutes to inactivate the enzyme and before being purified using a QAIquick nucleotide removal column (Qiagen), this was washed twice after binding to the column, and then eluted with 60 µL of nuclease free water to leave a 4 µM solution of radiolabelled S1*. PAGE Polyacrylamide gels were prepared by mixing 25 ml of 29:1 acrylamide/bis-acrylamide (National Diagnostic, protogel) with 5 mL of 10x Tris-Boric acid buffer (890 mM each, pH 8.3 or pH 7.05 adjusted with HCl) and 20 mL of Milli-Q water. To this 400 µL of a 10% w/v ammonium persulfate/water solution and 40 µL of TEMED were added to initialise polymerisation. This is then immediately poured between 2 glass plates and a comb inserted at the top this is then left to set for 1 hr. Samples were made up to 10 µl containing 0.4 µM per strand, of each strand (S1*, S2 and S3 unless stated otherwise), 1xTBN buffer (89 mM Tris, 89 mM Boric acid, 10 mM NaCl) and the indicated ratio of Complex or competitor. DNA, water, and buffer were mixed in solution before addition of the stated ratios of complex and competitor. Samples were then incubated at 37 degrees Celsius for 1 hr. 5 µl of 30% w/v glycerol was then added to each sample and the sample was then centrifuged, mixed, and pipetted into the wells on the gel. The gel was run at 140 V for 2 hours in 1xTB buffer. The gel was then removed from the plates and placed into a phosphor imaging box with a screen and left for 2 hours, the screen was then removed and imaged on a BIO-RAD FarosFX Plus molecular imager.

**Agarose gel electrophoresis:** Agarose gels were prepared by mixing 4 g of agarose (UltraPure™ Agarose, thermofisher scientific) with 400 ml of Milli-Q water (18.2 MΩ) containing 1x Tris-Boric acid buffer (890 mM each, pH 8.3). This was microwaved till all the solid had dissolved and cast into the agarose gel tray with a 15-lane comb. DNA/complex samples were prepared in 1xTB buffer with 30 µM DNA in base pairs and the labelled ratio of complex to this. Samples were 20 µl in total, to this, 10 µl of 30% w/v glycerol solution was added for sample loading into the gel. A DNA ladder was used in the left and right most lanes with 10, 4 and 0.5 kDa mass, top to bottom. The Gel was run at 140 V for 2 hours in 1xTB buffer. The gel was then removed from the plates and stained using SYBR™ Gold Nucleic Acid Gel Stain (Thermofisher scientifc) in 1xTB buffer for 45 minutes before imaging on a bio-rad ChemiDoc fluorescent imager with 305 nm excitation.

**pBR322 plasmid linearization preparation:** pBR322 plasmid DNA was purchased from MERCK, linearisation was achieved using the pst1-HF restriction endonuclease (new England biolabs). 20 µl of 0.5 µg/µl pBR-322 DNA solution was incubated with 10 µl of pst1-HF (20,000 units/ml), 50 µl of 10x cutsmart buffer and 420 µl Milli-Q water (18.2 MΩ) were incubated together at 37 ºC for 1 hr. This reaction mixture was then purified using QAIquick PCR purification columns and eluted with 50 µl Milli-Q water (18.2 MΩ). Concentrations in DNA base pairs was determined using a multichannel nanodrop 8000 spectrophotometer by using the absorbance at 260 nm and ε260 = 13,200 $mol^{-1}$ $dm^3$ $cm^{-1}$ using the beer lambert law. Confirmation of the linearization was confirmed using agarose gel electrophoresis by observing the characteristic band shift and by comparison to the 4 kDa band in the DNA ladder.

**Antiproliferative assays** Human lung adenocarcinoma (A549) cells were obtained from American Type Culture Collection (ATCC). The cells were cultured in Dulbecco's Modified Eagle Medium (DMEM, 4.5 g/L glucose Corning), supplemented with 10% fetal bovine serum (FBS, One shot, Gibco) and 1% penicillin/streptomycin (Gibco) and incubated in a humidified atmosphere of 5% $CO_2$ at 37 °C. To evaluate the antiproliferative effects of the compounds, cells were seeded in 96-well plates (Costar, Corning) at a concentration of 8000 cells/well and grown for 24 h in 200 µL culture medium. Solutions of the pillarplexes with the required concentration (1 to 100 µM) were prepared by diluting a freshly prepared stock solution ($10^{-2}$ M in MilliQ) of the corresponding compound in aqueous DMEM medium, accordingly. After 24 h of incubation, 200 µL of the compounds' dilutions in DMEM medium were added to each well and the cells were incubated for 24 h or 72 h, under standard culture conditions. Afterwards, the medium was replaced with a 3-(4,5-dimethyl-2-thiazoyl)-2,5-diphenyltetrazolium bromide (MTT, Fluorochem) solution in 10x PBS (Corning) at a final concentration of 0.5 mg/mL and incubated for 3-4 h. Following incubation, the MTT solution was aspirated from the wells and the purple formazan crystals were dissolved in DMSO. Absorbance at 550 nm was determined in quadruplicates for each condition, using a multi-well plate spectrophotometer (Victor X5, Perkin Elmer). Cell viability was determined as the ratio of absorbance between treated and untreated cells (untreated controls). The $EC_{50}$ value was calculated as the concentration showing a 50% reduction in cell viability, when compared to untreated controls, using a nonlinear fit of cell viability vs dose with GraphPad Prism 7 software. Data is represented as mean ± SEM of at least three independent experiments.

**Cell uptake ICP-MS analysis** For metal uptake/accumulation studies, cells were seeded in T25 flasks (Corning), grown to approximately 50% confluency and incubated with the corresponding metallodrug (dissolved directly in the incubation medium) at 6 µM for 24 h. At the end of the incubation period, cells were washed three times with ice cold PBS (containing 10 mM phosphate) before being detached using an enzyme free dissociation solution (Millipore, UK). The cells were counted after trypan blue staining in a haemocytometer, and the final suspension kept for analysis on the whole cell metal content. For whole cell extract analysis, cells were pelleted for 5 min at 700 x g and 4 °C and washed once with ice cold PBS. Cell lysis was achieved using a freeze-thaw technique suitable for cell uptake studies. Cold RIPA buffer (ThermoFisher Scientific) at a concentration of 1 mL per 5 million cells was added, followed by 30 sec of sonification with 50% pulse. After 15 min incubation on ice, the mixture was centrifuged at 14.000 xg at 4 °C for 15 minutes to pellet debris. Supernatant was transferred to a new Eppendorf and both were frozen at -80 °C. All samples were analyzed for their protein content prior to ICP-MS determination. For cell fractionation studies, the cell suspension following compound's treatment and PBS washing steps was divided into two equal aliquots. One aliquot was pelleted and kept for analysis on the whole cell, while the second aliquot was treated for cell fractionation as described below. For cell fractionation, the Nuclear/Cytosol Extraction Kit (Biovision Inc.) was used. All samples were digested in concentrated nitric acid for 3 h and filled to a total volume of 8 ml with a mixture HCl/water. Indium was added as an internal standard at a concentration of 0.5 ppb. The metal content was quantitated by inductively coupled plasma mass spectrometry (ICP-MS), using an ICP-MS Agilent 7900 (Agilent Technologies, Cheshire, UK) instrument, equipped with an internal autosampler and a nebulizer at a sample uptake rate of 0.25 ml/min. The instrument was calibrated on a daily basis. ICP-MS parameters were: RF power 1560 W, dwell time 0.3 s, replicates 10, monitored isotopes $^{197}$Au and $^{109}$Ag. Au and Ag standards for ICP-MS measurements were derived from PlasmaCAL standards (SCP Science, Quebec, Canada). The Agilent MassHunter software package was used for data processing. The obtained results are the average ± SE of at least three experiments.

**Compounds:** Gold and silver Pillarplexes and the nickel and iron Cylinders $[M_2L_3][Cl]_4$ were prepared as previously described in:

Altmann, P.J.; Pöthig, A. Pillarplexes: A Metal–Organic Class of Supramolecular Hosts. *J. Am. Chem. Soc.* **2016**, *138* (40), 13171–13174.

Hannon, M.J.; Painting, C.L.; Jackson, A.; Hamblin, J.; Errington, W. An inexpensive approach to supramolecular architecture. *Chem. Commun.* **1997**, 1807–1808.

Kerckhoffs, J.M.C.A.; Peberdy, J.C.; Meistermann, I.; Childs, LJ.; Isaac, C.J.; Pearmund, C.R.; Reudegger, V.; Khalid, S.; Alcock, N.W.; Hannon M.J.; Rodger, A. Enantiomeric resolution of supramolecular helicates with different surface topographies. *Dalton Trans.*, **2007**, 734-742

### Molecular Dynamics Simulations

**Parameterisation of the gold pillarplex and iron(II) cylinder** Parameters and topology files for the gold pillarplex were created using MCPB.py [Ref 1] of AmberTools20 [Ref 2] and moved to GROMACS topology using parmed (https://github.com/ParmEd/ParmEd). The iron(II) cylinder was parameterised as detailed in [ref 5]

**Parameterisation of DNA** B-DNA was created with random sequence using NAB Ambertools [Ref 2]. The 3WJ structure was taken from the 3I1D PDB crystal structure after removing the helicate. The 4WJ structure was taken from the 1XNS PDB crystal structure after removal of the peptides and water molecules, and each arm of the DNA structure was shortened by 9 base pairs to reduce the computational cost of modelling the 4WJ. [Ref 3] All DNA was parameterised using the AMBER forcefield parmbsc1. [Ref 4]

**Molecular Dynamics Simulations** In all simulations of the B-DNA and 3WJ, DNA was placed with the pillarplex (or cylinder) in a dodecahedral box with periodic boundary conditions. In all simulations of the 4WJ, DNA was placed with the pillarplex in a cubic box with periodic boundary conditions. All systems were solvated using the TIP3P model and neutralised with $Na^+$ ions. Additional $Na^+$ and $Cl^-$ ions were added to reach a NaCl concentration of 50mM. Energy minimisation and equilibration was carried out as previously described [Refs 5 and 6] in GROMACS. [Ref 7] All simulations used a 2 fs time step and run on the BlueBEAR cluster at U. Birmingham. After the simulations had finished, the trajectories were processed to remove periodic boundary conditions, translations and rotations.
